# Spike Antibody Fc Drives Protection from SARS-CoV-2 Challenge in Macaques

**DOI:** 10.64898/2026.03.13.711527

**Authors:** Caolann Brady, Melissa Govender, Jack Mellors, Tom Tipton, Karen Gooch, Adriana Tomic, Miles W. Carroll

## Abstract

A definitive correlate of protection (CoP) for SARS-CoV-2 has yet to be formally established. Previously, using data from a series of non-human primate vaccine challenge studies, we reported that neutralising antibodies (NAbs) are the strongest candidate for clinical protection against COVID-19 and that spike binding antibody is the strongest candidate CoP for viral burden post-challenge. In this study, we further characterised the protective binding antibody profile by analysing spike antibody-dependent complement deposition, FcγR binding, isotype and antibody glycosylation. Using the machine learning platform SIMON, we demonstrate that antibody-dependent complement deposition (ADCD) and FcγR binding are strong candidate co-correlates for each of the post-challenge outcomes; viral load and lung pathology. We found that spike antibody sialylation closely followed by FcγR2A, was the spike antibody feature with the strongest negative correlation with histopathology score. Spike antibody ADCD, FcγR binding, isotype and glycosylation significantly differed by immunisation regimen and sex, which demonstrates the heterogeneity of immune mechanisms induced by different immunisation platforms. We conclude that spike binding antibody, with the protective functional characteristics described herein, is a candidate CoP that captures both protection from severe clinical disease and protection against a high viral burden. These findings should be taken into consideration for future SARS-CoV-2 vaccine development.

## Introduction

Severe acute respiratory syndrome coronavirus 2 (SARS-CoV-2) spike antibodies remain a valuable measure of vaccine efficacy (1–5). We previously reported the predictive power of both spike binding and neutralising antibody (NAb) in a non-human primate (NHP) challenge model involving ninety rhesus and cynomolgus macaques (RhMs and CyMs), vaccinated with candidate DNA, mRNA or chimpanzee adenoviral (ChAd)-vectored vaccines (6–10) (**Supplementary Figure 1, Supplementary Table 1-3**). Using a systems immunology approach we found that NAbs were the strongest predictor of clinical protection (i.e. low lung histopathology score 6-8 days post-challenge), while spike binding antibody was the strongest predictor of lung viral load 6-8 days post-challenge and throat viral load post-challenge (11). However, beyond quantifying IgG spike binding and neutralisation, the features and functionality of ‘protective antibody’ that establish clinical and viral protection against SARS-CoV-2 have yet to be fully elucidated.

Antibodies contain both an antigen binding domain (Fab) and a fragment crystallisable (Fc) region. The Fc region binds to Fc receptors (FcR) expressed on innate cells. Antibody Fc interactions with FcγR3A/CD16 on natural killer (NK) cells results in antibody-dependent NK cell activation (ADNKA) and the release of perforins and granzymes; an effector function collectively referred to as antibody-dependent cellular cytotoxicity (ADCC). FcγR2A/CD32 activation on classical (CD14^+^CD16^-^) monocytes, in addition to FcγR3A/CD16 activation on non-classical (CD14^-^CD16^+^) and intermediate monocytes (CD14^+^CD16^+^), mediate antibody-dependent cellular phagocytosis (ADCP) of opsonised pathogen or pathogen-infected cells. FcγR2B and FcγR3B receptors are inhibitory and dampen the above innate cell responses when bound by Fc. Antibody-Dependent Complement Deposition (ADCD) refers to complement deposition (12). Antibody Fc binds C1q, the first complement protein in a cascade of complement convertase reactions, that may culminate in the formation of the membrane attack complex (MAC). The MAC induces cell death by forming pores in pathogen and pathogen-infected cell membranes. Although cell-based functional assay best capture the potency of antibody Fc induction of downstream cellular effects, FcγR-binding assay are now commonly used as a surrogate for cell-based functional assay (13–22), and are the more appropriate approach to both defining and measuring correlates of protection due to their relative ease and superior reproducibility.

The effector functions of SARS-CoV-2 antibodies, i.e. antibody with an Fc domain that can induce ADCC, ADCP or ADCD, has been widely reported (23–27). Research to-date has underscored the central role for the Fc in protection against SARS-CoV-2 variants of concern (VOCs). For example, NAb titres following BNT162b2 mRNA vaccination were found to be lower against Omicron (BA.2) versus Wuhan, however, FcγR2A and FcγR3A binding was largely unaffected and protection against severe COVID-19 was still achieved (15). Conversely, in a phase III study of NVX-CoV2373, eight out of ten vaccine breakthrough cases were infected with Alpha (B.1.1.7) VOC (28) which harbours the E484K mutation that weakens concurrent binding to antibody Fab and Fc domains, thus abrogating Fc effector function (29). Furthermore, therapeutic monoclonal antibody (mAb) S309 remained protective against Omicron (BA.1, BA.1.1 and BA.2) VOC in K18-hACE2 transgenic mice despite a reduction in neutralisation potency, unless the Fc of the mAb was mutated to abrogate FcγR binding, which resulted in a loss of efficacy (30). However, unpicking the importance of functional antibody, relative to the importance of NAb titres, during a SARS-CoV-2 infection homologous to the SARS-CoV-2 immunisation, is challenging, particularly given the heterogeneity in a population’s vaccination and SARS-CoV-2 exposure history. Therefore, our aim was to garner the data from pre-clinical NHP studies from 2020 to address this unknown.

Antibody isotype is known to correlate with FcγR binding and ADCD. In humans, SARS-CoV-2 mRNA vaccination induced an IgG1 dominant response, followed by IgG3, while IgG2 and IgG4 responses are rare and undetectable following a prime-boost, respectively (31–34). IgG4 titres increase over time and are boosted following an additional BNT162b2 dose or a natural SARS-CoV-2 infection, which was associated with a reduction in FcγR2A binding, ADCD and ADCP (33).

FcγR2A binding is largely dictated by antibody isotype. However, evidence suggests that FcγR3A binding is also highly dependent on IgG Fc glycosylation signatures, particularly N-linked glycosylation at N297 within the C_H_2-C_H_3 interface of the Fc domain. The core N297 glycan is central to IgG Fc functionality and N297 is highly preserved within the human species and between humans and macaques (35). Post-translational modifications (PTMs) of the N297 glycan core include core α1-6 fucosylation or branch terminal sialylation. Both fucosylation and sialylation are associated with reduced FcγR3A binding, while the absence of these residues is associated with increased FcγR3A binding. Sialylation is also associated with ADCD, and in the context of COVID-19, N297 sialylation increased ADCD by 3-4-fold and was associated with clinical protection (36, 37).

We screened ninety RhMs and CyMs from the NHP studies described in Brady *et al.* (11) for spike antibody isotype and IgG subclass, sialylation, FcγR binding and ADCD, using serum collected on the day of SARS-CoV-2 challenge following vaccination or prior challenge. Using a systems immunology approach with the machine learning platform SIMON, we aimed to assess how strongly these features associate with protection relative to previously identified spike-binding antibody titres and neutralising antibodies.

## Results

### Antibody Dependent Complement Deposition correlates with clinical protection

The finding that spike IgM was the strongest predictor of low lung viral load in Brady *et al.* (11), prompted analysis into the ADCD response, as IgM is the most potent activator of the classical complement pathway (38).

We found that ADCD did not significantly differ between any two vaccine groups across the study (Dunn’s multiple comparison test) (**Figure 1a**). In **Figure 1a** we also found ADCD responses to be inconsistent within groups. Low dose mRNA vaccinated macaques induced the most consistently low ADCD response, however, one macaque had a serum ADCD >100 complement activating units (CAU)/mL (standardised using an anti-SARS-CoV-2 antibody calibrant, see **Methods**). The re-challenge and FIV-vaccinated groups contained the animals with the greatest ADCD response, however there were also low responders in these groups, such that the mean ADCD response is similar across the one and two dose DNA, the FIV and the re-challenge groups.

**Figure 1.**
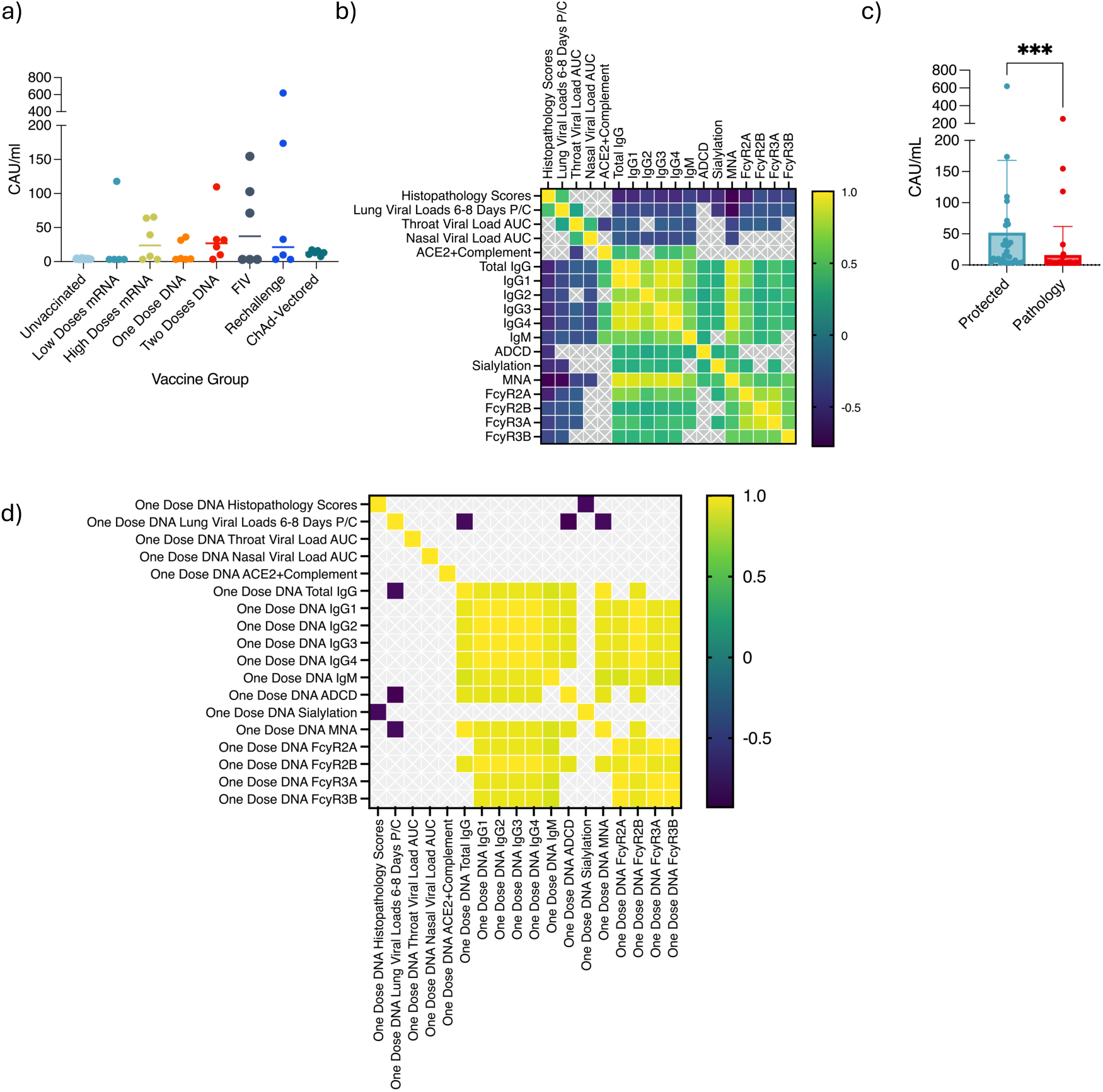
Antibody-Dependent Complement Deposition. **a)** ADCD by vaccine group. Datapoints represent individual macaques and the bars represent the mean. **b)** Heatmaps demonstrating all significant correlations between outcomes and spike antibody measurements when all data across studies is pooled (Two-sided Spearman correlation). **c)** ADCD comparison between clinically protected macaques and macaques with pathology. Two-sided Mann Whitney test performed. Datapoints represent individual macaques and the bars represent the mean and SD shown. **d)** Heatmaps demonstrating significant correlations between outcomes and spike antibody measurements when NHPs are analysed in the one dose DNA vaccine group (Two-sided Spearman correlation).

To investigate the relationship between ADCD and spike binding antibody titres across the isotypes and IgG subclasses measured previously (Brady et al. (11)), we performed correlation analysis. ADCD correlated significantly, but weakly with spike IgG titres measured by ELISA (r = 0.395, p = 0.0003) and bead assay (r = 0.357, p = 0.0007), spike IgG1 titres (r = 0.37, p = 0.0004), spike IgG2 titres (r = 0.266, p = 0.0129), spike IgG3 titres (r = 0.342, p = 0.0012), spike IgG4 titres (r = 0.365, p = 0.001) and spike IgM titres (r = 0.295, p = 0.006) (**Figure 1b**). When the correlation analysis was performed by group, only the one dose DNA vaccine group had a significant correlation between ADCD and IgG, IgG1, IgG2, IgG3, IgG4 titres (Spearman correlation, all r = 0.928, p = 0.022, **Figure 1d**).

To determine if ADCD was associated with protection post-challenge, further correlation analysis was performed. ADCD did not correlate with any viral load outcome, but ADCD did show a significant negative correlation with histopathology scores (r = -0.52, p < 0.0001, **Figure 1b**). This correlation between ADCD and histopathology scores aligns with the significant difference in ADCD observed between protected and pathology groups (Mann Whitney test; p = 0.0002, **Figure 1c**).

In a by-group analysis, within the mRNA, FIV and two dose DNA vaccine groups, there were no significant correlations between ADCD and post-challenge outcomes. In the one dose DNA vaccine group with low spike IgG titres, ADCD negatively correlated with lung viral load 6-8 days post-challenge (r = -0.928, p = 0.022, **Figure 1d**). Nasal viral load positively correlated with ADCD in the ChAd-vectored vaccine group (r = 0.943, p = 0.017, **Supplementary Figure 2**).

### FcγR Binding significantly differed by vaccine group and correlated with post-challenge outcomes

To ascertain whether FcγR binding, was associated with post-challenge outcome, spike antibody isotype and IgG subclass, and immunisation strategy, we measured anti-spike antibody binding to FcγR2A, FcγR2B, FcγR3A and FcγR3B.

Anti-spike antibody binding to FcγR2A, FcγR2B, FcγR3A and FcγR3B significantly differed by vaccine group (Kruskal Wallis test, p <0.0001, **Figure 2a-c and Supplementary Figure 3a**). As expected, vaccinated macaques had significantly higher FcγR binding compared to unvaccinated macaques. High dose mRNA, one and two dose DNA, FIV vaccination, and re-challenge macaques had significantly higher FcγR2A binding than unvaccinated macaques (Dunn’s multiple comparisons test, p < 0.0001, p = 0.0007, p = 0.0001, p = 0.0168, p = 0.0188, respectively, **Figure 2a**), High dose mRNA and one and two dose DNA vaccinated macaques also had significantly higher FcγR3A binding than unvaccinated macaques (Dunn’s multiple comparisons test, p < 0.0001, p = 0.0034 and p = 0.0013, respectively, **Figure 2b**) but also ChAd-vectored vaccinated macaques (Dunn’s multiple comparisons test, p = 0.004, p = 0.0418 and p = 0.023, respectively, **Figure 2b**). A similar trend was observed with the inhibitory FcγRs, with significantly lower FcγR2B and FcγR3B binding in unvaccinated and ChAd-vectored vaccinated macaque versus high dose mRNA and one and two dose DNA vaccinated macaques (**Figure 2c** and **Supplementary Figure 3a**). There were no further significant differences between vaccine groups.

**Figure 2.**
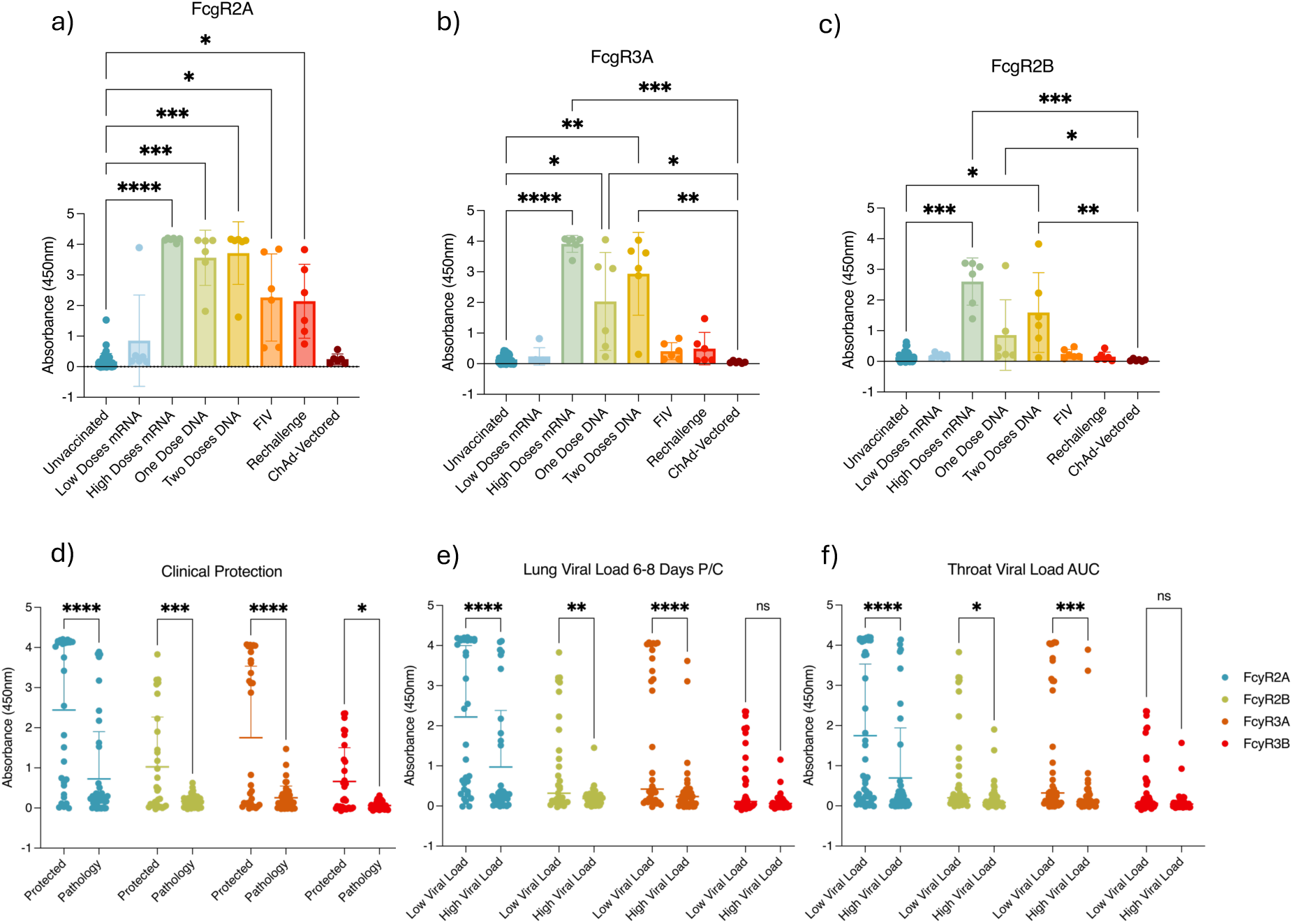
FcγR Binding. **a)** FcγR2A, **b)** FcγR3A, **c)** FcγR2B binding by vaccine group. Dunn’s multiple comparisons test performed. Datapoints represent individual macaques and the bars represent the mean and SD. FcγR binding comparison between post-challenge across groups; **d)** Clinical protection, **e)** Lung viral load 6-8 days post-challenge (P/C), **f)** Throat viral load area-under-the-curve (AUC). Tukey’s multiple comparisons test performed. Datapoints represent individual macaques and the bars represent the mean and SD.

With regards to clinical protection, FcγR binding had significant negative correlations with histopathology scores. FcγR2A most strongly correlated with histopathology score (r = -0.611, p < 0.0001) followed by FcγR3A (r = -0.431, p = 0.0004), while the correlations with FcγR2B and FcγR3B were weaker (r = -0.34, p = 0.006 and r = -0.384, p = 0.002, respectively). FcγR2A (p < 0.0001), FcγR2B (p = 0.0005), FcγR3A (p < 0.0001) and FcγR3B (p = 0.0139) binding was also significantly higher in macaques in the protected group versus the pathology group (Tukey’s multiple comparisons test, **Figure 2d**). The correlation between throat and lung viral load outcomes with each FcγR was weak (all r < 0.35, Figure **1b**), although FcγR2A, FcγR2B and FcγR3A binding was significantly higher in macaques in the low virus group versus the high virus group (Lung; p < 0.0001, p = 0.0074, p < 0.0001, respectively, and Throat; p < 0.0001, p = 0.0034 and p = 0.0002, respectively, Tukey’s multiple comparisons test, **Figure 2e and f**). FcγR binding did not significantly correlate with nasal viral load (**Figure 1b**). The only correlation between FcγR binding and outcomes in the analysis performed within vaccine groups was the significant negative correlation between FcγR2A and FcγR3A binding and nasal viral load AUC (r = -0.9429, p = 0.01667 for both, **Supplementary Figure 2**) in the high dose mRNA group.

The correlations between Ig isotypes and IgG subclasses with FcγR binding was stronger than with ADCD. FcγR2A binding correlated most strongly with spike Ig (total spike IgG; r = 0.695, p < 0.0001, IgG1; r = 0.674, p < 0.0001, IgG2; r = 0.489, p < 0.0001, IgG3; r = 0.602, p < 0.0001, IgG4; 0.597, p < 0.0001, IgM; r = 0.557, p < 0.0001, **Figure 1b**). FcγR3A binding also correlated with spike IgG subclass titres, although less strongly than FcγR2A (total spike IgG; r = 0.479, p < 0.0001, IgG1; r = 0.483, p < 0.0001, IgG2; r = 0.347, p < 0.0001, IgG3; r = 0.439, p < 0.0001, IgG4; 0.454, p < 0.0001, IgM; r = 0.439, p < 0.0001, **Figure 1b**). Inhibitory FcγRs weakly correlated with each spike Ig measurement (all r < 0.4, **Figure 1b**).

Stronger correlations between IgG subclasses and FcγR binding were observed within vaccine groups, most notably between FcγR2A binding and IgG1, IgG3, and IgG4 in the one dose and two dose DNA, FIV, and ChAd-vectored vaccine groups (all p < 0.035, r > 0.88, **Figure 1d** and **Supplementary Figure 1**). The one and two dose DNA vaccine group also showed correlations between IgG2 and FcγR2A binding (p < 0.035, r > 0.88, **Figure 1d** and **Supplementary Figure 2**). In addition, FcγR3A binding also correlated with IgG1-4 in the one-dose DNA vaccine group and with IgG1, IgG3 and IgG4 in the two dose DNA vaccine group (all r = 0.9429, p = 0.017, **Figure 1d** and **Supplementary Figure 2**).

### Spike Antibody Fc Glycosylation is associated with clinical and viral protection and impacts Fc effector functions

Since ADCD or FcγR could not be explained by Ig isotype or IgG subclass profile alone, we next examined the spike antibody glycosylation, focusing on sialylation. The glycosylation signature of spike-binding antibody was assessed using biotinylated lectins, which are plant-derived proteins that bind specific glycans (39, 40).

We found that in macaques that were clinically protected and had lower lung viral load 6-8 days post-challenge, spike antibody sialylation was significantly higher (p = 0.0008 and p = 0.0344, **Figure 3a**). Spike antibody sialylation also significantly correlated with histopathology scores (r = -0.55, p = 0.001) and lung viral load 6-8 days post-challenge (r = -0.501, p = 0.002, **Figure 1b**). Spike antibody sialylation differed significantly by immunisation group (Kruskal Wallis test, p = 0.0065, **Figure 3b**), with higher sialylation in macaques receiving the high dose mRNA vaccination compared with the low dose mRNA vaccination or the re-challenge macaques (Dunn’s multiple comparison test, p = 0.0019 and p = 0.0485, **Figure 3b**). The one dose DNA vaccine group with low spike IgG titres, also had a significant negative correlation between spike antibody sialylation and histopathology scores (r = -0.893, p = 0.042, **Figure 1d**).

**Figure 3.**
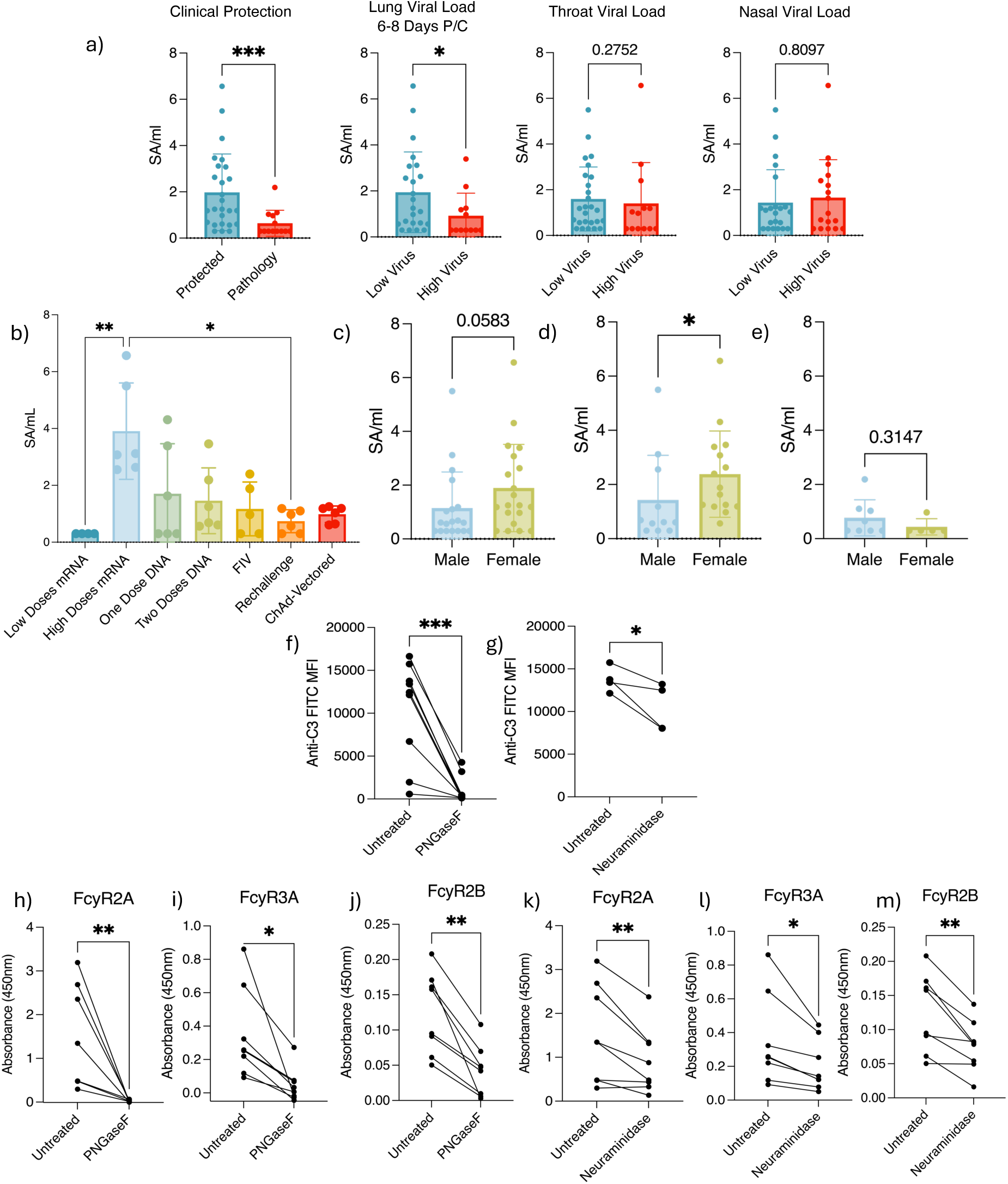
Spike Antibody Glycosylation Analysis. **a)** Spike antibody sialylation by post-challenge outcome - clinical protection, lung viral load 6-8 days post-challenge, throat viral load AUC and nasal viral load AUC. Datapoints represent each animal and the bars represent group means with SD. Two-sided Mann Whitney test performed. **b)** Spike antibody sialylation by immunisation group. Datapoints represent each animal and the bars show group means with SD. Non-parametric Kruskal Wallis test and Dunn’s multiple comparison test performed. **c)** Spike antibody sialylation by sex. Spike antibody sialylation by sex in all screened macaques, **d)** protected macaques and **e)** macaques with pathology. Two-sided Mann Whitney test performed. **f)** and **g)** ADCD responses in native versus deglycosylated serum. Datapoints represent the mean MFI of intra-assay technical ADCD replicates for macaque serum sample following **f)** PNGaseF treatment or **g)** neuraminidase-treatment, versus untreated. **g)** Includes the serum of macaques with high sialylation only (based on the measurement of spike antibody sialylation by the lectin-based assay). A two-tailed paired student t-test was performed with the biological replicates.

In female macaques, spike antibody sialylation trended higher compared to the male macaques, however, this was not significant (Two-sided Mann Whitney, **Figure 3c**). In clinically protected female macaques however, these animals had significantly higher spike antibody sialylation than protected male macaques (Two-sided Mann Whitney, p = 0.0315, **Figure 3d**). In macaques with pathology, there was no significant difference in spike antibody sialylation between sexes (Two-sided Mann Whitney, **Figure 3e**). This trend was not observed within all immunisation groups, and there were no significant differences in spike antibody sialylation between sexes within immunisation groups (Two-way ANOVA, Tukey’s multiple comparisons test), although samples sizes were low (**Supplementary Figure 5**).

Sialylation significantly correlated with ADCD (r = 0.3168, p = 0.0494), FcγR2A and FcγR3A binding (r = 0.3394, p = 0.0345 and r = 0.3484, p = 0.0297, respectively, Spearman correlation, **Figure 1b**). To determine the effect of antibody glycosylation and, specifically sialylation, on ADCD and FcγR binding, N-linked glycans and sialic acid residues were cleaved from test serum, using glycosidase enzymes, PNGaseF and neuraminidase, respectively. Four macaque serum samples with high sialylation and four samples with low sialylation were randomly selected for this analysis (**Supplementary Figure 6a**). Overnight treatment of macaque serum with PNGaseF resulted in a significant reduction in ADCD (Paired student t-test p = 0.0061, **Figure 3f**), FcγR2A (p = 0.0066), FcγR2B (p = 0.001) and FcγR3A (p = 0.0116) (Paired student t-test, **Figure 3h-j**). In macaques with high sialylation, neuraminidase treatment resulted in a significant decrease in ADCD (Paired student t test, p = 0.0485, **Figure 3g**). All macaque serum treated with neuraminidase treatment significantly reduced FcγR2A (p = 0.0093), FcγR2B (p = 0.0037) and FcγR3A (p = 0.0154) binding, irrespective of antibody sialylation level (**Figure 3k-m**).

Finally, to ensure there was no change in ADCD/FcγR binding signal due to improved/impaired antigen binding by IgG following deglycosylation, we assessed the ability of PNGaseF- and neuraminidase-treated serum to bind spike beads, and found no significant difference between the anti-IgG PE MFI of native versus PNGaseF-treated or neuraminidase-treated serum (**Supplementary Figure 6b**).

### Complement-Enhanced Neutralisation by Inhibition of ACE2 Binding

When investigating the relationship between ADCD and spike antibody data, the strongest correlation was observed between ADCD and NAb titres (Spearman correlation, r = 0.417, p = 0.0032, **Figure 1b**). When analysed by group, the correlation between ADCD and NAb titres was particularly strong in the one dose DNA vaccine groups (r = 0.928, p = 0.0022, **Figure 1d**).

Therefore, we investigated the possibility of complement-enhanced neutralisation, previously reported for SARS-CoV-2 (41) and other viruses (42). ACE2 inhibition assays were used as a surrogate for microneutralisation assays (MNA), to investigate the effect of complement on serum antibody-mediated neutralisation of spike ACE2 binding. We found that ACE2 inhibition significantly correlated with MNA data (Pearson correlation, r = 0.86, p < 0.0001, **Supplementary Figure 7**).

There was a significant increase in Wuhan spike ACE2 inhibition with non-HI complement relative to the HI complement condition (Paired student t test; p = 0.0003, n = 38, **Figure 4a**), suggesting complement-enhancement of ACE2 inhibition. The mean percentage difference from the HI complement condition was 14.17%. However, the response to the addition of complement was variable (95% CI: 1.366-26.97%) and there was a significant difference in complement-enhanced ACE2 inhibition between vaccine groups (One-way ANOVA; p = 0.0048, **Figure 4b**). The re-challenge group had significantly greater log_2_fold change in complement-enhanced ACE2 inhibition than the low dose mRNA group (Tukey’s multiple comparison test; p = 0.0104), the one dose DNA group (p = 0.0031) and the ChAd-vectored vaccine group (p = 0.0137) (**Figure 4b**). The mean percentage difference between complement and HI complement, may therefore be conflated by the rechallenge group, which had a mean percentage difference of 64.3%. The magnitude of complement-enhanced ACE2 inhibition was more homologous within vaccine groups, with the low dose mRNA group, the one dose DNA group, and the ChAd-vectored vaccine group being consistently low.

**Figure 4.**
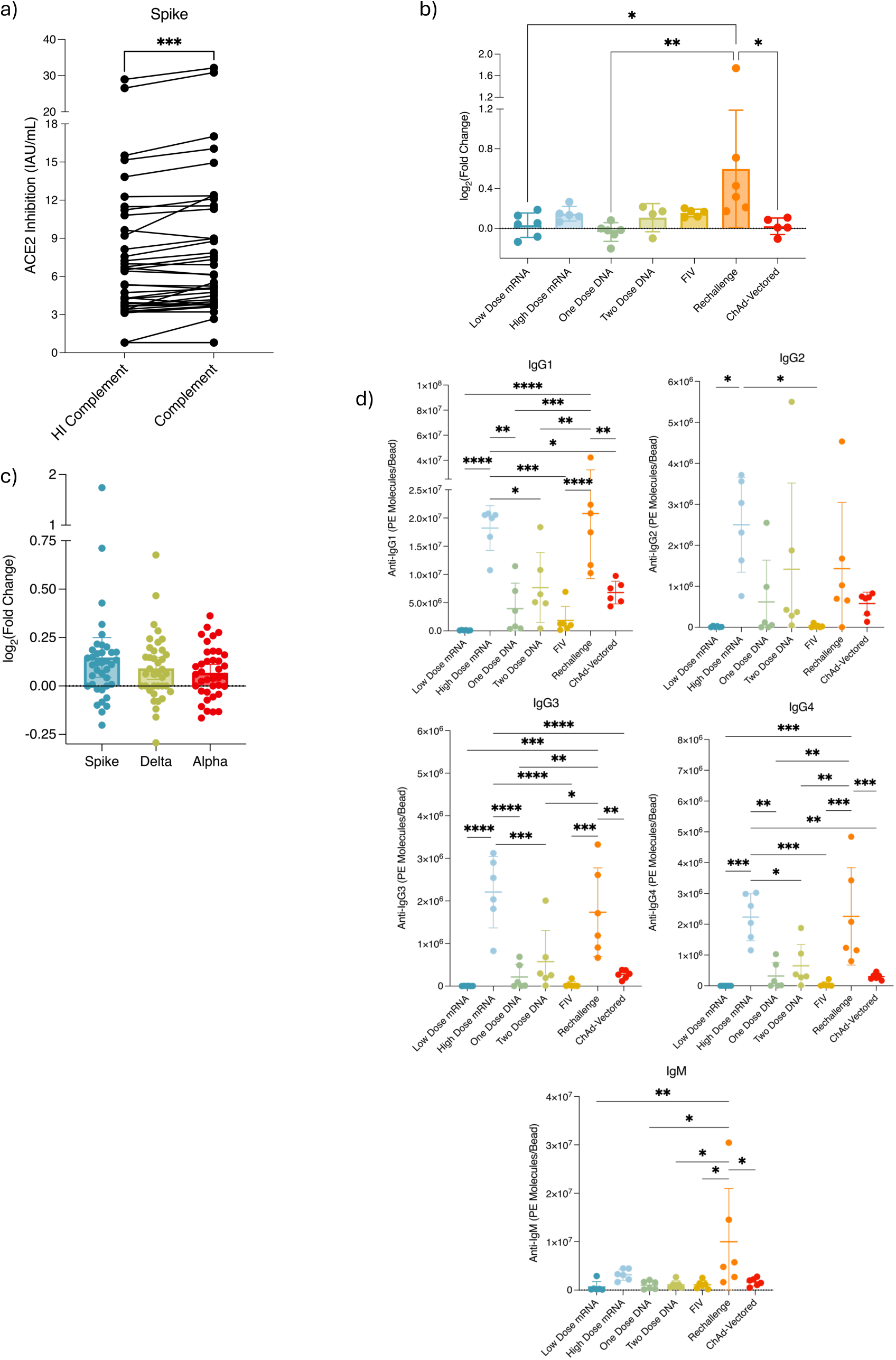
ACE2 Inhibition with Complement Analysis. **a)** Complement-enhancement of ACE2 Inhibition. Datapoints represent the mean ACE2 inhibition of technical replicates of SARS-CoV-2 immunised macaque serum ACE2 inhibition when heat-inactivated (HI)-complement versus non-HI complement is added to the ACE2 inhibition assay. Two-tailed paired student t-test performed. **b)** The log2(fold change) of when non-HI complement is added to the ACE2 inhibition assay versus HI-complement is added, by vaccine group. Datapoints represent the mean of technical replicates for each macaque tested. Tukey’s multiple comparisons test performed. **c)** The log2(fold change) in ACE2 inhibition by SARS-CoV-2 spike VOC. Datapoints represent mean of technical replicates for each macaque tested. **d)** IgG subclass and IgM by group. Datapoints represent mean of technical replicates for each macaque tested. Tukey’s multiple comparison test performed.

The log_2_fold complement-enhancement of ACE2 inhibition negatively correlated with throat viral load (r = -0.443, p = 0.006), but there were no further correlations between ACE2 inhibition with complement, and post-challenge outcomes (**Figure 1b**). There were no significant correlations within groups with log_2_fold complement-enhancement of ACE2 inhibition (data not shown).

The log_2_fold complement-enhancement of ACE2 inhibition correlated with total IgG and IgM titres (r = 0.429, p = 0.008 and r = 0.599, p < 0.0001, respectively), in addition to IgG1, IgG3 and IgG4 titre-ratio profiles (all r >0.449, p<0.005, **Figure 1b**). IgG1, IgG3 and IgG4 titres in both the rechallenge group and the high dose mRNA vaccine group were significantly higher than in the low dose mRNA, one dose DNA, two dose DNA, FIV and ChAd-vectored vaccine groups (all p < 0.036, Tukey’s multiple comparisons test, **Figure 4d**). Only high dose mRNA vaccine group IgG2 titres were significantly higher than any other group (low dose mRNA; p = 0.0118, and FIV; p = 0.0126) (**Figure 4d**). IgM titres in the rechallenge group only were also significantly higher than in the low dose mRNA, one dose DNA, two dose DNA, FIV and ChAd-vectored vaccine groups (all p < 0.027) (**Figure 4d**).

The addition of complement also significantly enhanced ACE2 inhibition of the Delta (B.1.617.2) and Alpha (B.1.1.7) variants (Paired student t test; p = 0.0088, p = 0.0209, respectively), however, the mean percentage difference from the HI complement condition was lower for these VOCs than for Wuhan spike (7.2%, 95% CI: 2.7-11.8%, and 5.3%, 95% CI: 1.6-9.7%) (**Figure 4c**). Lastly, complement did not significantly enhance Beta (B.1.351) ACE2 inhibition (B.1.351), and most samples were below the limit of detection for the assay (**Supplementary Figure 8**).

### SIMON Analysis

Examining the spike antibody Fc and its effector function across a range of vaccine platforms in the NHP pre-clinical model is important in the wider context of SARS-CoV-2 vaccine development. However, the extent to which the Fc predicts clinical and viral protection relative to spike binding or neutralising antibody, as previously reported in Brady *et al.* (11), remains unclear.

To address this, we used the Sequential Iterative Modelling Overnight (SIMON) platform to identify supervised machine learning (ML) algorithms that best model the post-challenge outcomes of clinical protection (i.e. low histopathology score) and viral loads in the lung 6-8 days post-challenge. For the purpose of ML analysis, the post-challenge outcomes were binarised; protection versus pathology for the clinical protection outcome, and low versus high virus for the lung viral load outcome, as described in the **Methods** and in Brady *et al.* (11). A total of 167 immune features measured at distinct matched timepoints were now included in the analysis; 158 immune parameters previously featured in Brady *et al.* (11)) and now also includes anti-spike antibody ADCD, FcγR binding, sialylation and complement-enhancement of ACE2 inhibition. Area under receiver operator curve (AUROC) was used to evaluate model performance, as described in the **Methods**.

Using SIMON, 171 ML algorithms were tested and 28 models successfully modelled clinical protection. The model with the strongest performance was built using the svmPoly algorithm (train AUROC = 0.8465, test AUROC = 0.9259, **Supplementary Figure 9a**), which performed better (by test AUROC) than the clinical protection model in Brady *et al.* (11) (**Figure 5c**). To ensure that the predictive power of the SIMON models was robust and not driven by class imbalance, we also evaluated performance using multiple metrics beyond AUROC. The optimal model for clinical protection, svmPoly, demonstrated high classification accuracy on the unseen test set, achieving a Sensitivity of 1.0 and Specificity of 0.83.

**Figure 5.**
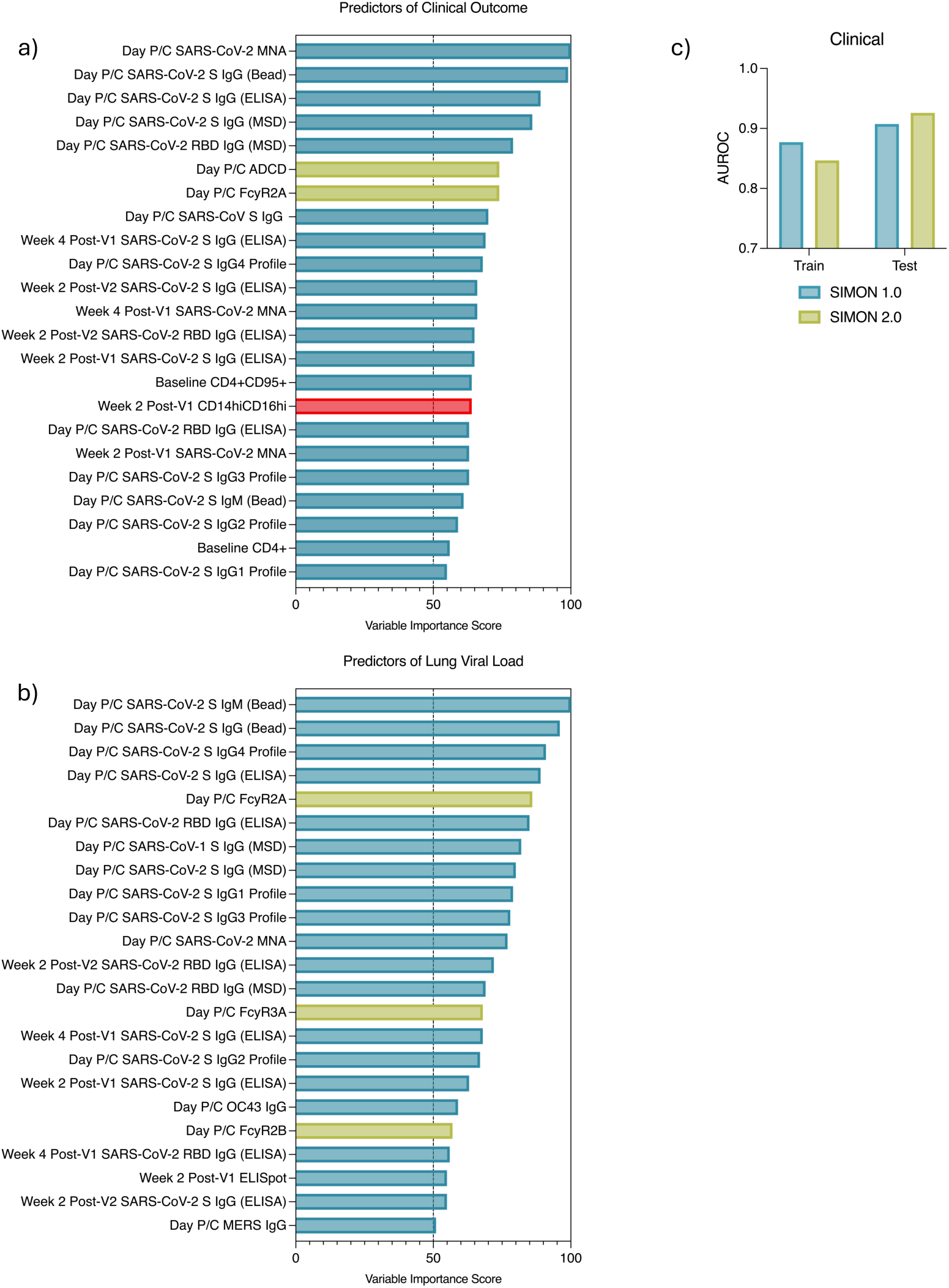
SIMON Analysis. SIMON outputs for identifying the immune predictors of **a)** Clinical Protection and **b)** Lung Viral Load 6-8 Days Post-Challenge. Graphs represent the Variables of Importance used by **a)** the svmPoly model to predict low (in blue) and high (in red) histopathology scores, **b)** the pcaNNet model to predict low (in blue) and high (in red) lung viral loads. Day P/C = sampled on the day of challenge, prior to challenge. Green represents the new variables of importance identified when functional antibody data was included in the modelling. **c)** Comparison between model performance from Brady *et al.* and with the dataset that now includes functional antibody data.

There were 23 immune features that were important for building the svmPoly model (variable importance score >50), and so these features best predicted clinical protection when modelled with the svmPoly algorithm (**Figure 5a**). 21 of 23 of these immune features, and their VIS scores, were shared between this svmPoly model with the functional antibody data, and the Boosted Classification Trees (ada) algorithm from Brady *et al*. (11). NAb titres remained the strongest predictor of protection from SARS-CoV-2 challenge with the highest VIS of 100. The new predictors which improved model performance (shown in green) included ADCD and FcγR2A binding on the day pre-challenge (both VIS = 74) (**Figure 5a**). ADCD and FcγR2A binding was significantly higher in protected versus pathology macaques as demonstrated above in **Figure 1c** and **Figure 2d** (p = 0.0002 and p < 0.0001, respectively). Although NAb titres remained the strongest predictor, the co-correlates - spike binding antibody, ADCD and FcγR2A binding - improved the predictive power of the model.

Nine models successfully modelled lung viral load 6-8 days post-challenge. The pcaNNet algorithm yielded the most balanced performance (Test AUROC = 0.78, Sensitivity = 0.71, Specificity = 0.83), identifying it as a more reliable predictor than algorithms with higher AUROC but poor specificity (**Supplementary Figure 9b**). There were 23 immune features that were important for building the pcaNNet model (**Figure 5b**). 20 of 23 of these immune features, and their VIS scores, were shared between this pcaNNet model with functional antibody data and the LogitBoost algorithm from Brady *et al*. (11). Spike IgM titres remaining as the strongest predictor of lung viral load 6-8 days post-SARS-CoV-2 challenge with the highest VIS of 100 (**Figure 5b**). FcγR2A (VIS 86), FcγR3A (VIS 68) and FcγR2B (VIS 57) were the three new predictors (shown in green), in line with the higher FcγR binding in the low lung virus versus high lung virus groups (FcγR2A; p < 0.0001, FcγR2B; p = 0.0074, FcγR3A; p < 0.0001, **Figure 2e**). Performance did not improve upon the lung viral load in Brady *et al.,* unlike for clinical protection above, therefore FcγR2A, FcγR3A and FcγR2B are likely only surrogates of protection against high viral burden in the LRT.

Throat viral load could not be robustly modelled, on the grounds of poor sensitivity and specificity. Nasal viral load could also not be modelled, on the grounds of poor AUROC performance, as was also the case in Brady *et al*. (11). This is in line with the observation that FcγR binding did not significantly differ between the low and high nasal virus groups (**Supplementary Figure 3b**) and there were no correlations between FcγR binding and nasal viral loads (**Figure 1b**).

## Discussion

Spike antibodies have been consistently associated with protection against SARS-CoV-2 in pre-clinical models and clinical studies (2, 5, 43–45). However, the precise mechanisms underlying this protection remain unclear. Is binding or neutralisation of the spike protein alone sufficient to prevent severe COVID-19, or does optimal humoral immunity also require Fc-mediated effector functions? If Fc engagement is indeed important, which antibody isotypes, IgG subclasses, or glycosylation patterns confer a functional Fc domain, and binding which FcRs are crucial for protection? To address these questions and extend our understanding of functional antibody responses in COVID-19 protection, we analysed a cohort of 90 NHPs, focusing on the relationship between spike-specific antibodies, their Fc properties, and protective outcomes.

Given that ADCD reflects the capacity of antibodies to activate complement and eliminate infected cells, we evaluated whether ADCD responses were associated with clinical protection from SARS-CoV-2. ADCD negatively correlated with histopathology score, distinguishing protected macaques from those with pathology, and was amongst the strongest predictors of clinical protection, further supporting its role in clinical protection from COVID-19 (29, 46–48). ADCD also correlated moderately with total spike IgG, spike IgM, and all IgG subclasses, consistent with the structural similarity of macaque IgG subclasses (49, 50). These moderate correlations, together with the variation in ADCD responses across vaccination groups, suggests that additional antibody features beyond subclass or isotype likely contribute to ADCD responses, which may be independent of the immunisation strategy.

Among humoral responses, NAb titres showed the strongest (albeit moderate) association with ADCD. To further explore this relationship, we investigated complement-mediated enhancement of neutralisation with the ACE2 competition assay, a surrogate assay for neutralisation (51). Addition of exogenous complement significantly increased ACE2 inhibition relative to HI complement. Complement also enhanced neutralisation of SARS-CoV-2 VOCs that the cohort of macaques had no prior exposure to. The vaccine group showing the greatest magnitude of effect was the rechallenge group, while the other vaccine groups exhibited significantly lower magnitudes. These observations are consistent with those by Mellors *et al.*, who reported cohort-dependent variability in complement-mediated enhancement (41). Overall, these findings further support the re-introduction of exogenous complement into neutralisation assays using HI test serum, to better reflect the true neutralisation potential of serum samples.

Using a lectin-based assay, we analysed spike antibody sialylation and found it significantly correlated with NAb titres, ADCD, and FcγR2A and FcγR3A binding. The association of sialylation with complement and NAb titres may reflect enhanced germinal centre antigen deposition by a mechanism that involves complement deposition on sialylated antibody (52). Quast *et al.* reported that sialic acid residues sterically hinder C1q binding (53), therefore desialylation may be expected to improve ADCD. We observed a modest reduction in ADCD and FcγR binding following removal of sialic acid residues by neuraminidase treatment, relative to more marked effect of PNGaseF cleavage of N-glycans. Therefore, rather than directly improving ADCD, sialylation may instead act as a marker of a more mature, high-affinity antibody response, as reflected in its correlation with NAb titres. Spike antibody sialylation negatively correlated with histopathology scores and lung viral load 6-8 days post-challenge and was significantly higher in protected macaques indicating a potential protective role for sialylation in modulating antibody function and limiting disease severity. The one dose DNA vaccine group, which had significantly lower spike IgG titres than the other vaccine groups (11), exhibited the strongest correlations of all of the vaccine groups between histopathology score and sialylation, and lung viral load 6-8 days post-challenge, MNA and ADCD (**Figure 1d** and **Supplementary Figure 2**). This suggests that even if antibody titres are low, antibody sialylation, ADCD and neutralisation can dictate protection. Antibody sialylation levels also varied by vaccine group and sex. Consistent with Wang *et al.* (32), spike antibody sialylation was significantly higher in macaques after two high dose mRNA vaccines compared with natural infection (re-challenge). Female macaques showed a trend toward higher sialylation, with clinically protected females exhibiting significantly higher levels than males, suggesting a sex-linked influence on antibody glycosylation and function.

Similar to ADCD and spike antibody sialylation, FcγR binding negatively correlated with histopathology scores. In particular, FcγR2A showed the strongest association with post-challenge outcomes, correlating not only with reduced histopathology scores and lung viral load at days 6-8 but also with lower throat viral load AUC. Incorporating FcγR2A into predictive models, alongside ADCD, significantly improved clinical protection model performance. Although NAbs remained the strongest overall predictor of clinical protection, these findings suggest that NAbs equipped with FcγR2A-binding and ADCD activity represent an optimal protective antibody profile. FcγR2A, together with FcγR2B and FcγR3A predicted low lung viral load, with FcγR2A outperforming NAb titres as an independent predictor. These results indicate that non-neutralising, but FcγR-binding antibodies contribute meaningfully to viral control and disease mitigation. While NAbs restrict viral entry, effective clearance still requires Fc-mediated mechanisms. FcγR2A binding, commonly used as a surrogate for ADCP, reflects monocyte-mediated uptake and apoptotic elimination of opsonised virus without promoting inflammation, thereby reducing pathology and viral load (13–22). Similarly, FcγR3A binding, serves as a surrogate for NK cell-mediated ADNKA and ADCC (13–22). Our findings indicate that FcγR binding plays an important role in limiting viral load and pathology, at least in the lower respiratory tract, but cell-based functional assays would be required to prove downstream functionality. FcγR binding responses within and between vaccine groups was however, heterogenous, therefore FcγR binding may not be a vaccine-specific CoP.

Negative correlations between post-challenge outcomes and each of the FcγRs, suggest that both higher inhibitory FcγR2B and higher activating FcγR2A binding are protective. Because macaque activating and inhibitory receptors share more similar affinities than human receptors (50, 54, 55), the net effect of FcγR2A and FcγR2B binding is uncertain without cell-based functional assays where the intracellular signalling domains dictate activation or inhibition. However, analysis of the FcγR2A to FcγR2B binding ratios across vaccination groups revealed a more balanced FcγR2A-FcγR2B binding profile in some groups and relatively higher FcγR2A binding in others, including the rechallenge group, which had significantly higher FcγR2A:2B ratios than the mRNA vaccine groups (**Supplementary Figure 10**). Therefore, at a minimum, we can say that the macaque spike antibody profile, induced by different immunisation strategies result in different FcγR2A:FcγR2B binding ratios.

FcγR2A binding strongly correlated with spike-specific IgG1, IgG3, IgG4, and IgM titres, while IgG2 titres showed only moderate correlation with FcγR2A. FcγR3A exhibited moderate correlation with spike-specific isotypes, while FcγR2B correlations with spike isotypes were generally weak. Correlations between ADCD and antibody isotypes were weaker than those observed for FcγR2A and FcγR3A binding with these antibody isotypes. Among all IgG subclasses, IgG2 titres also consistently correlated most weakly with FcγRs. Additionally, IgG2 was also the only isotype that did not correlate with complement-enhancement of ACE2 inhibition, and IgG2 was typically a poorer predictor of clinical protection and lung viral load compared with the other IgG subclasses. These observations suggest that IgG2 behaves similarly across human and RhM and CyM macaque species, as the relatively least ‘functional’ IgG subclass. Notably, macaque IgG2 binds FcγR3A and FcγR2B more strongly than human IgG2 (50), suggesting that correlations with this subclass might have been stronger had macaque FcγR proteins been used. In contrast, IgG1 correlated most strongly with the FcγRs, FcγR2A, FcγR2B and FcγR3A, consistent with macaque IgG1 functioning like human IgG3. However, IgG1 titres were the least predictive of the IgG subclasses of clinical outcome ie. lung pathology, in the SIMON analysis.

Interestingly, higher IgG4 titres and ratios emerged as the most protective subclass, strongly associated with reduced pathology and lower lung viral loads. In humans, IgG4 responses are rare and correlate with reduced FcγR2A binding, ADCD and ADCP (33), as to be expected with its anti-inflammatory phenotype. Unlike human IgG4, RhMs and CyMs lack a destabilising hinge residue required for Fab-arm exchange (FAE)(49, 56), that makes it more reactive than its human counterpart, exhibiting higher affinity for FcγR3A (50). This interspecies variation in IgG4, likely underlies IgG4’s strong correlations with ADCD and FcγR binding in this study, highlighting IgG4 as a key contributor to protective effector functions in macaques. This is likely a macaque-specific finding.

All four IgG subclasses predicted clinical protection and lower lung viral load, indicating that class-switching and a co-ordinated multi-subclass humoral response is important for controlling SARS-CoV-2 in macaques. In fatal human COVID-19 cases, the cause of death is not typically driven by viraemia, but rather by multi-organ failure resulting from a cytokine storm (57, 58). In contrast, SARS-CoV-2-challenged naïve macaques typically develop only subclinical to mild disease and resolve infection by day 15-20 (8), indicating that macaques effectively balance viral control with regulation of inflammation. This suggests that sufficient regulatory and anti-inflammatory mechanisms are important for protection against COVID-19. Consistent with this, higher antibody sialylation, which has anti-inflammatory properties, was associated with clinical protection in our study, whereas low sialylation has been linked to severe COVID-19 in human (36, 59). Species-specific differences between macaque and human immune systems along the activation-inhibition axis likely contribute to this regulatory capacity. Macaques exhibit higher baseline antibody sialylation, increased affinity of IgG2, and the absence of an extended hinge region in macaque IgG3, which in humans confers potent pro-inflammatory activity (60). Collectively, these features may render macaques less susceptible to hyperinflammation, as observed in *B.malayi* infection studies, where reduced IgG sialylation in macaques was required to initiate a robust immune response (61).

Several limitations to this study should be considered. First, the small sample sizes may have reduced statistical power in the analysis performed by vaccine group. Second, FcγR binding was assessed using human rather than macaque FcγR proteins, despite the known species differences in FcγR affinity (50, 54, 55). Consequently, the use of macaque FcγR may have yielded stronger correlations with macaque IgG subclasses. Third, both SNA and neuraminidase specificity for terminal sialic acid residues and cross-reactivity is uncertain, however, the minimal impact of neuraminidase on ADCD in low-sialylation samples suggests acceptable reagent specificity. Specificity of SNA for terminal acid residues also was confirmed by comparing SNA binding when treated or left untreated with neuraminidase enzyme which cleaves terminal sialic acid residues (**Supplementary Figure 4**). As we did not purify Ig isotypes to measure both sialylation and ADCD response post-neuraminidase treatment, we cannot confirm the sialylation signature of IgG or IgM or how these isotypes rely on sialylation for ADCD. Notably, Haslund *et al.* reported that desialylation of spike-specific IgM substantially reduced ADCD by 50%, but there was no substantial effect of desialylation on spike ADCD following neuraminidase treatment of plasma that had not been fractionated by isotype (62). In our dataset, a significant difference in spike ADCD following neuraminidase treatment was observed, but the effect was minimal. This underscores the need for isotype-specific glycosylation and functionality studies in macaques. Spike antibody sialylation did not correlate with IgM titres, therefore isotype-specific analyses will be essential for clarifying these relationships.

In our analysis of ninety SARS-CoV-2-challenged RhMs and CyMs, we have demonstrated immunisation platform-dependent and -independent variation in spike-specific humoral profiles. We have shown that certain features of this profile, i.e., FcγR binding, isotype and IgG subclass and ADCD, can predict clinical protection and viral load. Our analysis suggests that effective immunisation depends on generating B cell clones that produce antibodies capable of both neutralisation and C1q and FcγR binding with complement further enhancing neutralisation through deposition via the antibody Fc. No single IgG subclass uniquely defined functional activity in macaques, however, IgG4 emerged as the subclass most predictive of post-challenge outcomes in our ML analysis. The association of IgG4 and spike antibody sialylation with post-challenge outcomes, spike-specific ADCD and FcγR binding, suggests that macaques can achieve potent, yet regulated anti-inflammatory humoral responses that limit immunopathology during SARS-CoV-2 infection. In conclusion, NAbs remain a strong candidate CoP for protection against lung pathology, however, predictive accuracy of vaccine efficacy improves when complement deposition and FcγR binding are considered. Furthermore, our data suggests that serum spike antibody FcγR binding is a better predictor of lung viral burden than serum NAb titres.

## Methods

### Pre-Clinical Study Design

Ninety rhesus macaques (RhMs) and cynomolgous macaques (CyMs) were assigned to receive either a primary viral challenge, one or two doses of a vaccine candidate (including an mRNA vaccine, a DNA vaccine, a formalin-inactivated vaccine FIV), or a ChAdOx-vectored vaccine; see **Supplementary Table 1**) or to serve as unimmunised controls. Animals were subsequently challenged or re-challenged with a high dose of SARS-CoV-2 (5 × 10⁶ PFU of Victoria/1/2020) administered via intratracheal and intranasal routes (**Supplementary Figure 1**). In all groups except the FIV cohort, challenge occurred 28 days after the final or only vaccine dose; FIV-vaccinated animals were challenged 14 days post-vaccination. In the re-challenge cohort, a 28-day interval was maintained between the primary and secondary exposures.

During the acute phase of infection, nasal and throat swabs were collected for quantitative PCR (qPCR) analysis. Six-to-eight days after challenge, lungs were harvested for histopathological evaluation, and viral genomic RNA (gRNA) levels in bronchoalveolar lavage (BAL) fluid were measured by qPCR to assess lung viral load. In the ChAdOx and RhM-versus-CyM studies, animals euthanised at earlier or later timepoints were excluded from analyses of clinical outcomes and lung viral load (see **Supplementary Table 2** for group numbers). Lung pathology was graded according to the scoring system described by Salguero *et al.* (8), which evaluates seven parameters on a scale from 0 to 4, where 4 indicates more severe pathology, yielding a maximum cumulative score of 28 across all parameters.

All experimental work was conducted under the authority of a U.K. Home Office–approved project license (PDC57C033) that had been subject to local ethical review at UK Health Security Agency Porton Down by the Animal Welfare and Ethical Review Body.

### Antibody-Dependent Complement Deposition

This protocol is based on ADCD assay described in (42, 63, 64). Spike protein was covalently coupled to carboxylated paramagnetic beads (Spherotech #CMPAK-4068-6K) as described previously (11). Test serum was heat inactivated (HI) at 56°C for 30min and diluted in blocking buffer with spike-conjugated magnetic beads. Following a 2-hour incubation, plates were washed twice with wash buffer. A 10% complement solution was prepared with IgG/IgM depleted human complement serum (Pelfreez #34010-1) and incubated with beads for 30 minutes at 37°C. Following a wash step, FITC-conjugated rabbit anti-human C3c polyclonal antibody (Abcam #ab4212), was incubated with beads for 20 minutes in the dark. The well contents were resuspended in HBSS for flow cytometry analysis using the BD LSRFortessa™X-20 Cell Analyser. Complement activation units (CAU) was assigned to each test sample and was calculated using a 12-point standard curve of the Anti-SARS-CoV-2 Antibody Diagnostic Calibrant (20/162 NIBSC, UK). The FITC MFI of blank samples were subtracted as background FITC fluorescence prior to interpolation from the standard curve.

### MSD ACE2 inhibition and complement assay

The ability of antibodies to inhibit SARS-CoV-2 spike (Wuhan, Delta B.1.617.2, Alpha B.1.1.7 and Beta B.1.351) binding to ACE2 was assessed in macaque serum samples with the V-PLEX SARS-CoV-2 Panel 13 ACE2 Kit (Meso Scale Diagnostics, K15466U-2). The assay was performed as per manufacturer’s instructions and as previously described (41). Briefly, wells were blocked for 30 minutes and washed, before the addition of test and control (human COVID-19 positive and negative) sera, and the ACE2 Calibration Reagent 3 for 1 hour with either 20% complement or heat-inactivated complement. After incubation, SULFO-TAG Human ACE2 Protein was added for an additional 1-hour incubation with the samples. After washing, MSD GOLD Read Buffer B was added, and the plates were read with the MESO QuickPlex SQ 120MM instrument. The raw data was analysed with MSD Discovery Workbench software v4, with standard curves created by fitting the signals from the standard using a 4-parameter logistic model. Inhibition was presented as IAU/ml and graphs created with Graphpad Prism 9.

### SARS-CoV-2 FcγR ELISA

Ninety-six well plates were coated with 5 µg/mL spike protein (LakePharma #46328) in DPBS either overnight at 4°C or for 2 hours at 37 °C. All remaining steps were performed at room temperature (RT). Wells were washed with 0.05% Tween-PBS and blocked with blocking buffer for 30 minutes. After a subsequent washing step, test sera and controls (human COVID-19 positive and negative) were diluted in blocking buffer and added to the plate for 1 hour. Complexes of 1µg/ml of each human FcγR protein: CD32A/FcγR2A, CD32B/FcγR2B, CD16A/FcγR3A, CD16B/FcγR3B (2B Scientific Limited), were diluted in HRP-streptavidin (1:2000 in blocking buffer, 0.5 mg/ml) and incubated on a roller for 10 minutes. After sample removal and washing, each FcγR-HRP-Streptavidin complex was added to wells for 30 minutes. Wells were washed once more, and 100 µl TMB substrate was added for 10-15 minutes and incubated in the dark. To stop the reaction, 100 µl/well of TMB stop was added and the absorbance was read 450 nm. Data was presented as absorbance and graphed using Graphpad Prism 9.

### Sialylation

This protocol was based on the whole IgG SNA binding ELISA described in (40). Spike-conjugated APC fluorescent carboxyl magnetic beads (50beads/μL), were incubated with a 100AU/mL of serum (diluted as per calculated neat concentration by ELISA, as described in (11)) at 25 °C for 2 hours. During the incubation, biotinylated SNA (140kDa, VectorLabs #B-1305-2) and streptavidin-PE (BioLegend#405203) conjugates were prepared in a 4:1 molar ratio (as recommended in (65)), and a quarter of the total streptavidin-PE volume to be added, was added to the mix every 5 minutes, for a total incubation time of 30min. The bead-antibody complexes were washed with HBSS using the EasySep™ Magnet. Streptavidin-PE-SNA conjugates, diluted to 1μg/mL in HBSS, were added for 1.5 hours at 37°C. The beads were washed and resuspended in HBSS for flow cytometry.

### Enzyme Treatment

Serum was treated with PNGaseF (New England Biolabs #P0704) as per the manufacturer’s instructions. PNGaseF was diluted 1:11 with 1μL of test sera and 1X Glycobuffer 2, and incubated overnight at 37°C. For 2-3,6,8 neuraminidase (New England Biolabs #P0720) treatment was diluted 1:6 with 1μL of test sera and 1X Glycobuffer 1 and incubated overnight at 37°C in a total reaction volume of 12μL.

### SIMON Analysis

The dataset was formatted such that each parameter at each timepoint was put forward as a candidate immune predictor of protection for SIMON analysis. Data pre-processing was performed by SIMON which included data centring and scaling, imputation of missing values (medianImpute) and removal of features with zero or near-zero variance.

Depending on the outcome of interest, a different number of macaques were included in each analysis. For example, animals that were not culled between the timeframe of 6-8 days post-SARS-CoV-2 challenge and did not have a CT scan performed during acute infection, were excluded from the clinical protection analysis (n=12). Animals that were not culled between the timeframe of 6-8 days post-SARS-CoV-2 challenge (n=20) or did not have post-challenge lung outcomes assessed (n=1) could not be included in the lung viral load analysis. The median was used as the cut-off to define binary outcomes.

Pathology scores, used to determine clinical protection outcomes, were assigned to SARS-CoV-2-challenged lungs harvested post-cull by veterinary pathologist using the published scoring system (8). In circumstances where pathology scoring was unavailable for the macaque, CT scores during acute COVID-19 were used to assess protection status (occurred in 14 cases). Macaques with pathology scores above the median pathology score (median=6.75) were less protected, and represented the ‘pathology’ group, whilst those below the median pathology score represented the ‘protected’ group. The median appropriately divided the animals across the groups (**Supplementary Figure 11**). The outcome of viral protection in the nose, throat and lung, was based on the quantification of SARS-CoV-2 nucleocapsid RNA by RT-qPCR in nasal washes, throat swabs and bronchiolar lavage (BAL) fluid respectively. Area under the curve (AUC) of the nasal and throat PCR results were calculated to broadly capture viral control at these sites during acute infection. The median of nasal PCR AUC (median=1.5 ×10^7^), throat PCR AUCs (median=590,000) and BAL PCR from 6-8 days post-challenge (median=120,000) defined the cut-off for the high and low virus groups. The median appropriately divided the animals across the groups (**Supplementary Figure 12, 13, 14**). Macaques that lacked PCR data were excluded from the analysis (nine macaques for lung viral load analysis, none for nasal or throat viral load analysis). The density plots were generated using R statistical software v4.1.2.

Data was partitioned 80% in the training set and 20% in the testing set. Model performance was summarised by receiver operator curves (ROC) of sensitivity versus the false positive rate (1-specificity). Area under ROC (AUROC) was used as the metric of model performance. Model acceptance criteria was a train AUROC>0.7, and a test AUROC>train AUROC, so as not to select an overfitted model. The magnitude of the contribution of immune features to the building of the models was reported as a VIS within a range of 0-100. Immune features scoring >50 are VOIs and represent candidate predictors of protection.

## Acknowledgements

We would like to express our gratitude to the UKHSA for their support and valuable contributions in the original supply of NHP samples, and to Professor Tess Lambe and her laboratory for providing reagents and guidance for the initial experiments used to optimise the FcγR assay.

This work was funded by the Coalition of Epidemic Preparedness Innovations (CEPI), under grant number H5R01920 awarded to Professor Miles Carroll (M.C.). The funders did not contribute to or influence paper conceptualisation, data analysis or interpretation, or writing of the manuscript.

## Contributors

Conceptualization: M.C.

Methodology: M.C., A.T., C.B., T.T., J.M., M.G.

Data Curation and Investigation: C.B. and M.G.

Data Analysis, Interpretation, Visualisation: C.B.

Funding acquisition: M.C.

Project administration: K.G., T.T., M.C.

Supervision: T.T, A.T., M.C.

Writing – original draft: C.B.

Writing – review & editing: All authors reviewed and commented on the manuscript.

## Data Availability

The raw data for all of immune predictors identified by the SIMON analysis in Brady et al. 2025 (DOI: 10.1038/s41541-025-01103-2) was provided in the Supplementary Appendices. Any additional data will be made available on reasonable request to the corresponding author.

## Bibliography

1. Earle KA, Ambrosino DM, Fiore-Gartland A, Goldblatt D, Gilbert PB, Siber GR, et al. Evidence for antibody as a protective correlate for COVID-19 vaccines. Vaccine. 2021;39(32):4423–8.

2. Khoury DS, Cromer D, Reynaldi A, Schlub TE, Wheatley AK, Juno JA, et al. Neutralizing antibody levels are highly predictive of immune protection from symptomatic SARS-CoV-2 infection. Nature Medicine. 2021;27(7):1205–11.

3. Benkeser D, Fong Y, Janes HE, Kelly EJ, Hirsch I, Sproule S, et al. Immune correlates analysis of a phase 3 trial of the AZD1222 (ChAdOx1 nCoV-19) vaccine. npj Vaccines. 2023;8(1):36.

4. Goldblatt D, Fiore-Gartland A, Johnson M, Hunt A, Bengt C, Zavadska D, et al. Towards a population-based threshold of protection for COVID-19 vaccines. Vaccine. 2022;40(2):306–15.

5. Gilbert PB, Montefiori DC, McDermott AB, Fong Y, Benkeser D, Deng W, et al. Immune correlates analysis of the mRNA-1273 COVID-19 vaccine efficacy clinical trial. Science. 2022;375(6576):43–50.

6. Bewley KR, Gooch K, Thomas KM, Longet S, Wiblin N, Hunter L, et al. Immunological and pathological outcomes of SARS-CoV-2 challenge following formalin-inactivated vaccine in ferrets and rhesus macaques. Science advances. 2021;7(37):eabg7996-eabg.

7. Gooch KE, Smith TRF, Salguero FJ, Fotheringham SA, Watson RJ, Dennis MJ, et al. One or two dose regimen of the SARS-CoV-2 synthetic DNA vaccine INO-4800 protects against respiratory tract disease burden in nonhuman primate challenge model. Vaccine. 2021;39(34):4885–94.

8. Salguero FJ, White AD, Slack GS, Fotheringham SA, Bewley KR, Gooch KE, et al. Comparison of rhesus and cynomolgus macaques as an infection model for COVID-19. Nature Communications. 2021;12(1):1260.

9. Rauch S, Gooch K, Hall Y, Salguero FJ, Dennis MJ, Gleeson FV, et al. mRNA vaccine CVnCoV protects non-human primates from SARS-CoV-2 challenge infection. bioRxiv. 2020:2020.12.23.424138.

10. Lambe T, Spencer AJ, Thomas KM, Gooch KE, Thomas S, White AD, et al. ChAdOx1 nCoV-19 protection against SARS-CoV-2 in rhesus macaque and ferret challenge models. Commun Biol. 2021;4(1):915.

11. Brady C, Tipton T, Carnell O, Longet S, Gooch K, Hall Y, et al. A systems biology approach to define SARS-CoV-2 correlates of protection. npj Vaccines. 2025;10(1):69.

12. Zhang A, Stacey HD, D’Agostino MR, Tugg Y, Marzok A, Miller MS. Beyond neutralization: Fc-dependent antibody effector functions in SARS-CoV-2 infection. Nature Reviews Immunology. 2023;23(6):381–96.

13. Kaplonek P, Cizmeci D, Fischinger S, Collier A-r, Suscovich T, Linde C, et al. mRNA-1273 and BNT162b2 COVID-19 vaccines elicit antibodies with differences in Fc-mediated effector functions. Science Translational Medicine. 2022;0(0):eabm2311.

14. Boudreau CM, Burke JS, Yousif AS, Sangesland M, Jastrzebski S, Verschoor C, et al. Antibody-mediated NK cell activation as a correlate of immunity against influenza infection. Nature Communications. 2023;14(1):5170.

15. Haycroft ER, Davis SK, Ramanathan P, Lopez E, Purcell RA, Tan LL, et al. Antibody Fc-binding profiles and ACE2 affinity to SARS-CoV-2 RBD variants. Med Microbiol Immunol. 2023;212(4):291–305.

16. Selva KJ, Ramanathan P, Haycroft ER, Reynaldi A, Cromer D, Tan CW, et al. Preexisting immunity restricts mucosal antibody recognition of SARS-CoV-2 and Fc profiles during breakthrough infections. JCI Insight. 2023;8(18).

17. Bartsch YC, Tong X, Kang J, Avendaño MJ, Serrano EF, García-Salum T, et al. Omicron variant Spike-specific antibody binding and Fc activity are preserved in recipients of mRNA or inactivated COVID-19 vaccines. Sci Transl Med. 2022;14(642):eabn9243.

18. Kaplonek P, Cizmeci D, Kwatra G, Izu A, Lee JS-L, Bertera HL, et al. ChAdOx1 nCoV-19 (AZD1222) vaccine-induced Fc receptor binding tracks with differential susceptibility to COVID-19. Nature Immunology. 2023;24(7):1161–72.

19. Tong X, McNamara RP, Avendaño MJ, Serrano EF, García-Salum T, Pardo-Roa C, et al. Waning and boosting of antibody Fc-effector functions upon SARS-CoV-2 vaccination. Nature Communications. 2023;14(1):4174.

20. Kaplonek P, Fischinger S, Cizmeci D, Bartsch YC, Kang J, Burke JS, et al. mRNA-1273 vaccine-induced antibodies maintain Fc effector functions across SARS-CoV-2 variants of concern. Immunity. 2022;55(2):355–65.e4.

21. Bartsch YC, Cizmeci D, Kang J, Zohar T, Periasamy S, Mehta N, et al. Antibody effector functions are associated with protection from respiratory syncytial virus. Cell. 2022;185(26):4873–86.e10.

22. Adams LE, Leist SR, Dinnon KH, West A, Gully KL, Anderson EJ, et al. Fc-mediated pan-sarbecovirus protection after alphavirus vector vaccination. Cell Reports. 2023;42(4):112326.

23. Herman JD, Wang C, Loos C, Yoon H, Rivera J, Eugenia Dieterle M, et al. Functional convalescent plasma antibodies and pre-infusion titers shape the early severe COVID-19 immune response. Nature Communications. 2021;12(1):6853.

24. Natarajan H, Crowley Andrew R, Butler Savannah E, Xu S, Weiner Joshua A, Bloch Evan M, et al. Markers of Polyfunctional SARS-CoV-2 Antibodies in Convalescent Plasma. mBio. 2021;12(2):10.1128/mbio.00765-21.

25. Bégin P, Callum J, Jamula E, Cook R, Heddle NM, Tinmouth A, et al. Convalescent plasma for hospitalized patients with COVID-19: an open-label, randomized controlled trial. Nature Medicine. 2021;27(11):2012–24.

26. Zohar T, Loos C, Fischinger S, Atyeo C, Wang C, Slein MD, et al. Compromised Humoral Functional Evolution Tracks with SARS-CoV-2 Mortality. Cell. 2020;183(6):1508–19.e12.

27. Bates TA, Lu P, Kang YJ, Schoen D, Thornton M, McBride SK, et al. BNT162b2-induced neutralizing and non-neutralizing antibody functions against SARS-CoV-2 diminish with age. Cell Reports. 2022;41(4):111544.

28. Heath PT, Galiza EP, Baxter DN, Bocito M, Browne D, Burns F, et al. Safety and Ecicacy of NVX-CoV2373 Covid-19 Vaccine. New England Journal of Medicine. 2021;385(13):1172–83.

29. Gorman MJ, Patel N, Guebre-Xabier M, Zhu AL, Atyeo C, Pullen KM, et al. Fab and Fc contribute to maximal protection against SARS-CoV-2 following NVX-CoV2373 subunit vaccine with Matrix-M vaccination. Cell Reports Medicine. 2021;2(9):100405.

30. Case JB, Mackin S, Errico JM, Chong Z, Madden EA, Whitener B, et al. Resilience of S309 and AZD7442 monoclonal antibody treatments against infection by SARS-CoV-2 Omicron lineage strains. Nature Communications. 2022;13(1):3824.

31. Farkash I, Feferman T, Cohen-Saban N, Avraham Y, Morgenstern D, Mayuni G, et al. Anti-SARS-CoV-2 antibodies elicited by COVID-19 mRNA vaccine exhibit a unique glycosylation pattern. Cell Reports. 2021;37(11):110114.

32. Chakraborty S, Gonzalez JC, Sievers BL, Mallajosyula V, Chakraborty S, Dubey M, et al. Early non-neutralizing, afucosylated antibody responses are associated with COVID-19 severity. Science Translational Medicine.14(635):eabm7853.

33. Irrgang P, Gerling J, Kocher K, Lapuente D, Steininger P, Habenicht K, et al. Class switch toward noninflammatory, spike-specific IgG4 antibodies after repeated SARS-CoV-2 mRNA vaccination. Science Immunology. 2023;8(79):eade2798.

34. Tejedor Vaquero S, de Campos-Mata L, Ramada JM, Díaz P, Navarro-Barriuso J, Ribas-Llaurado C, et al. The mRNA-1273 Vaccine Induces Cross-Variant Antibody Responses to SARS-CoV-2 With Distinct Profiles in Individuals With or Without Pre-Existing Immunity. Front Immunol. 2021;12:737083.

35. Crowley AR, Ackerman ME. Mind the Gap: How Interspecies Variability in IgG and Its Receptors May Complicate Comparisons of Human and Non-human Primate Effector Function. Frontiers in Immunology. 2019;10(697).

36. Larsen MD, de Graaf EL, Sonneveld ME, Plomp HR, Nouta J, Hoepel W, et al. Afucosylated IgG characterizes enveloped viral responses and correlates with COVID-19 severity. Science. 2021;371(6532):eabc8378.

37. Dekkers G, Treffers L, Plomp R, Bentlage AEH, de Boer M, Koeleman CAM, et al. Decoding the Human Immunoglobulin G-Glycan Repertoire Reveals a Spectrum of Fc-Receptor- and Complement-Mediated-Effector Activities. Front Immunol. 2017;8:877.

38. Cooper NR. The Classical Complement Pathway: Activation and Regulation of the First Complement Component11Publication number 3541 IMM. In: Dixon FJ, editor. Advances in Immunology. 37: Academic Press; 1985. p. 151–216.

39. Van Damme EJM. Lectins as Tools to Select for Glycosylated Proteins. In: Gevaert K, Vandekerckhove J, editors. Gel-Free Proteomics: Methods and Protocols. Totowa, NJ: Humana Press; 2011. p. 289–97.

40. Sołkiewicz K, Krotkiewski H, Jędryka M, Kratz EM. Variability of serum IgG sialylation and galactosylation degree in women with advanced endometriosis. Sci Rep. 2021;11(1):5586.

41. Mellors J, Dhaliwal R, Longet S, Tipton T, McInnes I, Siebert S, et al. Complement-mediated enhancement of SARS-CoV-2 antibody neutralisation potency in vaccinated individuals. Nature Communications. 2025;16(1):2666.

42. Mellors J, Tipton T, Fehling SK, Akoi Bore J, Koundouno FR, Hall Y, et al. Complement-Mediated Neutralisation Identified in Ebola Virus Disease Survivor Plasma: Implications for Protection and Pathogenesis. Frontiers in immunology. 2022;13.

43. Feng S, Phillips DJ, White T, Sayal H, Aley PK, Bibi S, et al. Correlates of protection against symptomatic and asymptomatic SARS-CoV-2 infection. Nature Medicine. 2021;27(11):2032–40.

44. Wei J, Matthews PC, Stoesser N, Newton JN, Diamond I, Studley R, et al. Protection against SARS-CoV-2 Omicron BA.4/5 variant following booster vaccination or breakthrough infection in the UK. Nature Communications. 2023;14(1):2799.

45. Wei J, Pouwels KB, Stoesser N, Matthews PC, Diamond I, Studley R, et al. Antibody responses and correlates of protection in the general population after two doses of the ChAdOx1 or BNT162b2 vaccines. Nat Med. 2022;28(5):1072–82.

46. McMahan K, Yu J, Mercado NB, Loos C, Tostanoski LH, Chandrashekar A, et al. Correlates of protection against SARS-CoV-2 in rhesus macaques. Nature. 2021;590(7847):630–4.

47. Yu J, Tostanoski LH, Peter L, Mercado NB, McMahan K, Mahrokhian SH, et al. DNA vaccine protection against SARS-CoV-2 in rhesus macaques. Science. 2020;369(6505):806–11.

48. Routhu NK, Gangadhara S, Lai L, Davis-Gardner ME, Floyd K, Shiferaw A, et al. A modified vaccinia Ankara vaccine expressing spike and nucleocapsid protects rhesus macaques against SARS-CoV-2 Delta infection. Science Immunology. 2022;7(72):eabo0226.

49. Tolbert WD, Subedi GP, Gohain N, Lewis GK, Patel KR, Barb AW, et al. From Rhesus macaque to human: structural evolutionary pathways for immunoglobulin G subclasses. MAbs. 2019;11(4):709–24.

50. Warncke M, Calzascia T, Coulot M, Balke N, Touil R, Kolbinger F, et al. Different adaptations of IgG effector function in human and nonhuman primates and implications for therapeutic antibody treatment. J Immunol. 2012;188(9):4405–11.

51. Tipton T, Laidlaw S, Nguyen D, Longet S, Barnes E, Carroll MW. ACE2 Inhibition ELISA Is an Effective Surrogate for SARS-CoV-2 Live Virus Neutralisation. Available at SSRN: https://ssrncom/abstract=5525615 or http://dxdoiorg/102139/ssrn5525615. 2025.

52. Lofano G, Gorman MJ, Yousif AS, Yu W-H, Fox JM, Dugast A-S, et al. Antigen-specific antibody Fc glycosylation enhances humoral immunity via the recruitment of complement. Science Immunology. 2018;3(26):eaat7796.

53. Quast I, Keller CW, Maurer MA, Giddens JP, Tackenberg B, Wang LX, et al. Sialylation of IgG Fc domain impairs complement-dependent cytotoxicity. J Clin Invest. 2015;125(11):4160–70.

54. Bruhns P, Iannascoli B, England P, Mancardi DA, Fernandez N, Jorieux S, et al. Specificity and affinity of human Fcγ receptors and their polymorphic variants for human IgG subclasses. Blood. 2009;113(16):3716–25.

55. Boesch AW, Miles AR, Chan YN, Osei-Owusu NY, Ackerman ME. IgG Fc variant cross-reactivity between human and rhesus macaque FcγRs. MAbs. 2017;9(3):455–65.

56. Labrijn AF, Rispens T, Meesters J, Rose RJ, den Bleker TH, Loverix S, et al. Species-specific determinants in the IgG CH3 domain enable Fab-arm exchange by acecting the noncovalent CH3-CH3 interaction strength. J Immunol. 2011;187(6):3238–46.

57. Ragab D, Salah Eldin H, Taeimah M, Khattab R, Salem R. The COVID-19 Cytokine Storm; What We Know So Far. Front Immunol. 2020;11:1446.

58. Tang Y, Liu J, Zhang D, Xu Z, Ji J, Wen C. Cytokine Storm in COVID-19: The Current Evidence and Treatment Strategies. Frontiers in Immunology. 2020;Volume 11 - 2020.

59. Chakraborty S, Gonzalez J, Edwards K, Mallajosyula V, Buzzanco AS, Sherwood R, et al. Proinflammatory IgG Fc structures in patients with severe COVID-19. Nature Immunology. 2021;22(1):67–73.

60. Boesch AW, Osei-Owusu NY, Crowley AR, Chu TH, Chan YN, Weiner JA, et al. Biophysical and Functional Characterization of Rhesus Macaque IgG Subclasses. Front Immunol. 2016;7:589.

61. Petralia LMC, Santha E, Behrens A-J, Nguyen DL, Ganatra MB, Taron CH, et al. Alteration of rhesus macaque serum N-glycome during infection with the human parasitic filarial nematode Brugia malayi. Scientific Reports. 2022;12(1):15763.

62. Haslund-Gourley BS, Woloszczuk K, Hou J, Connors J, Cusimano G, Bell M, et al. IgM N-glycosylation correlates with COVID-19 severity and rate of complement deposition. Nature Communications. 2024;15(1):404.

63. Barrett JR, Belij-Rammerstorfer S, Dold C, Ewer KJ, Folegatti PM, Gilbride C, et al. Phase 1/2 trial of SARS-CoV-2 vaccine ChAdOx1 nCoV-19 with a booster dose induces multifunctional antibody responses. Nature Medicine. 2021;27(2):279–88.

64. Tomic A, Skelly DT, Ogbe A, O’Connor D, Pace M, Adland E, et al. Divergent trajectories of antiviral memory after SARS-CoV-2 infection. Nature Communications. 2022;13(1):1251.

65. Weskamm LM, Dahlke C, Addo MM. Flow cytometric protocol to characterize human memory B cells directed against SARS-CoV-2 spike protein antigens. STAR Protoc. 2022;3(4):101902.

