## Supplementary Tables and Figures for "Spike Antibody Fc Drives Protection from SARS-CoV-2 Challenge in Macaques": Supplementary Figures_Final.pdf

| Manufacturer | Platform | Adjuvant | Antigen | Delivery | Citation |
|---|---|---|---|---|---|
| **Oxford-UKHSA** | Formalin-inactivated virus | Alhydrogel | Whole SARS-CoV-2 | Intramuscular | (6) |
| **Inovio** | DNA | Unknown | Full-length WT spike | Intradermal with CELLECTRA-ID electroporation technology | (7) |
| **Oxford-AstraZeneca** | ChAd-vector | Adenoviral vector | Full-length spike | Intramuscular | (9) |
| **CureVac** | Non-nucleoside modified mRNA | LNP | Pre-fusion stabilised full-length spike | Intramuscular | (10) |

**Supplementary Table 1.** Manufacturer, vaccine platform, adjuvant, antigen and delivery mechanisms of the vaccines included in this study.

| | Unvaccinated | Low Dose mRNA | High Dose mRNA | One Dose DNA | Two Dose DNA | FIV | ChAd-Vectored | Re-Challenge |
|---|---|---|---|---|---|---|---|---|
| **Clinical Outcome** | 35 | 6 | 6 | 6 | 6 | 6 | 6 | 6 |
| **LRT Viral Load** | 31 | 6 | 6 | 6 | 6 | 6 | 2 | 6 |

**Supplementary Table 2.** Number of macaques in each subgroup of study for each post-challenge outcome investigate

| Response | Vaccine Group Hierarchy of Median Response |
|---|---|
| **Spike IgG** | Rechallenge > High Dose mRNA > Two Dose DNA > ChAd-Vectored > One Dose DNA > FIV |
| **MNA** | High Dose mRNA > Two Dose DNA > ChAd-Vectored > One Dose DNA > FIV |
| **ELISpot** | FIV > ChAd-Vectored > High Dose mRNA > One/Two Dose DNA/Low Dose mRNA |
| **Histopathology Score** | Two Dose DNA < High Dose mRNA < One Dose DNA < FIV/ChAd-Vectored < Rechallenge < Low Dose mRNA |
| **Lung Viral Load** | High Dose mRNA < ChAd-Vectored < Two Dose DNA/Rechallenge < FIV < One Dose DNA |

**Supplementary Table 3.** Summary of findings by vaccine group, in Brady et al. 2025.

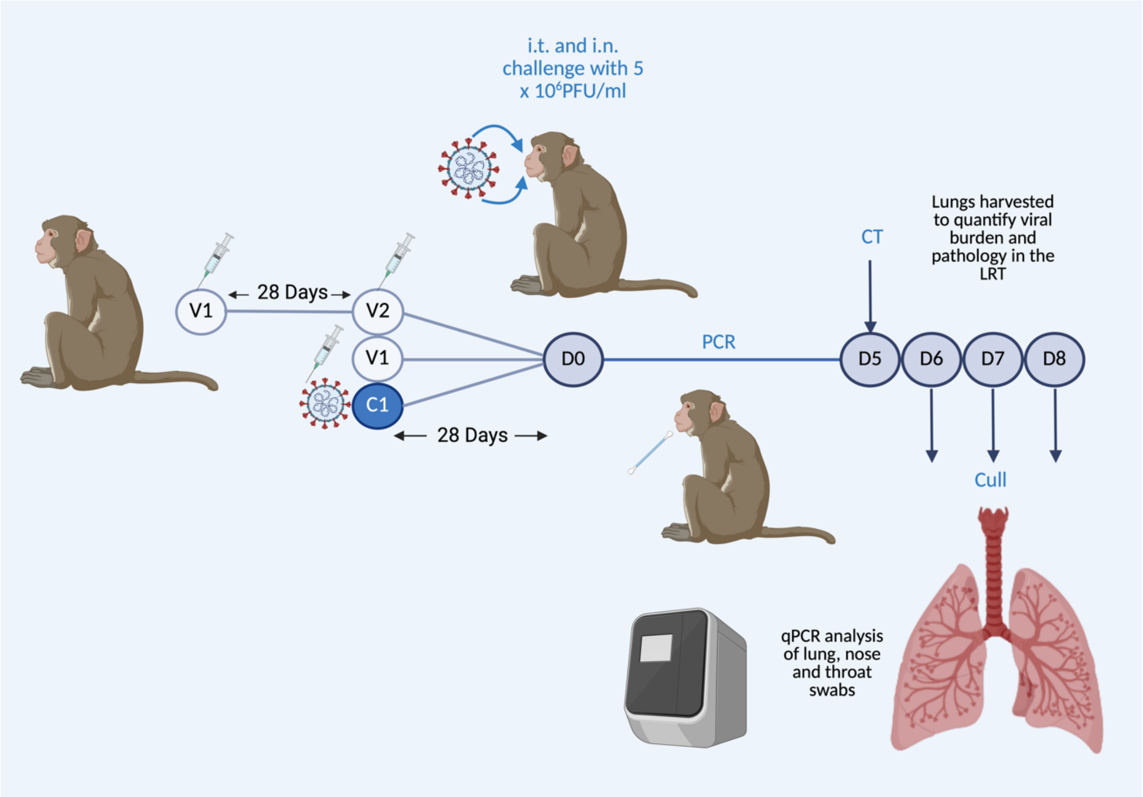


**Supplementary Figure 1. Summary of the study design and the primary analysis in Brady *et al.* 2025.** The day of viral challenge following immunisation was designated as D0. In two-dose regimens (mRNA and two dose DNA), a 28-day interval separated the first (V1) and second (V2) vaccine doses. A 28-day interval was also maintained between the second, or only, vaccine dose, and viral challenge, except for the FIV group, which was challenged 14 days after FIV vaccination. In the re-challenged cohort, there was a 28-day interval between the primary challenge (C1) and the secondary challenge (D0). During acute infection, nasal and throat swabs were collected for qPCR analysis. Animals were euthanised 6-8 days post-challenge (D6-D8), and lung tissues were harvested for histopathology analysis and scoring, which underpinned the categorizing of animals into ‘pathology’ and ’protected’ groups for the outcome of ’clinical protection;’. When comparing responses and outcomes by vaccine group, we observed the following results, as reported in Brady *et al.* 2025. *Figure adapted from Brady et al. 2025.*

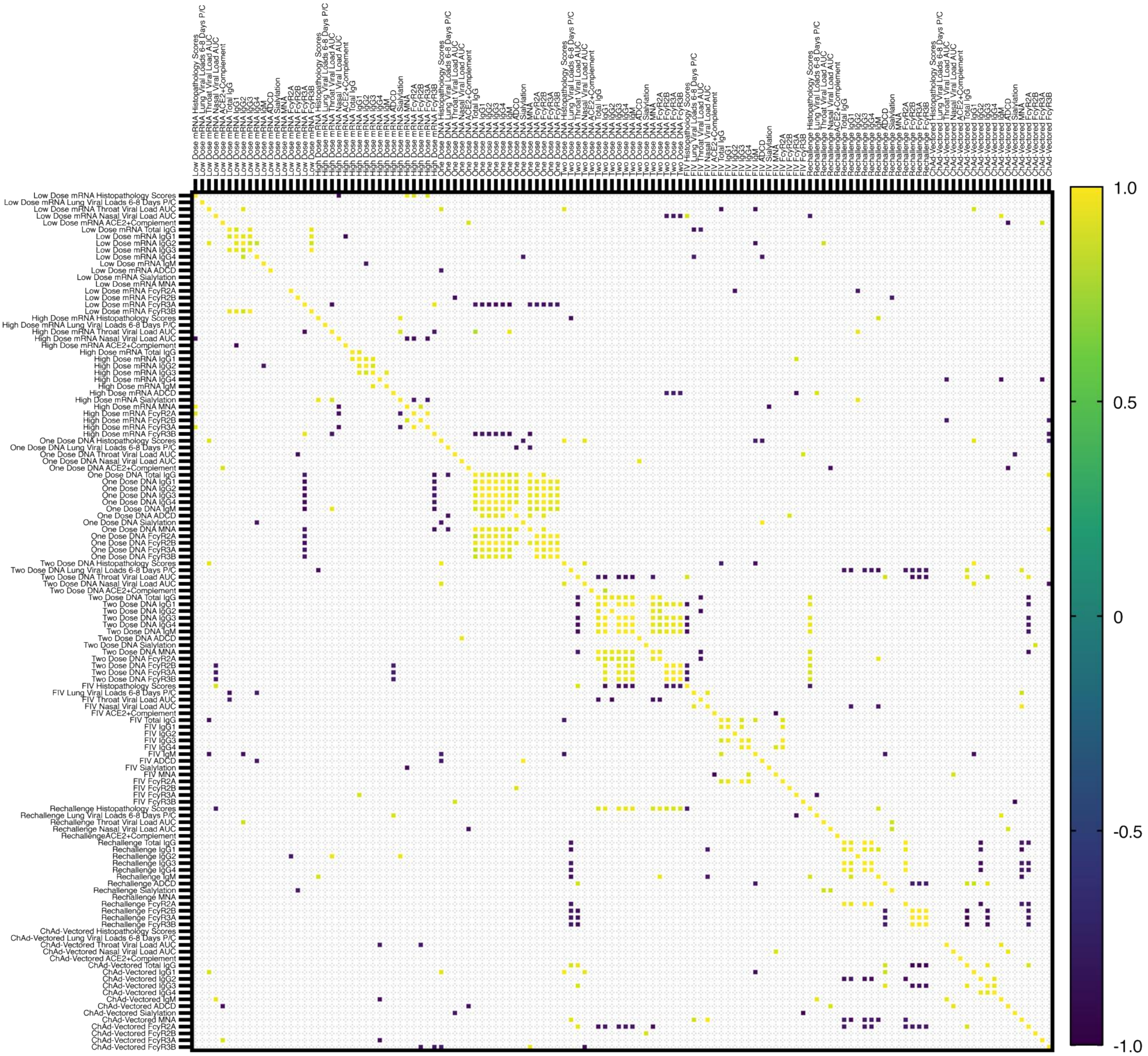


**Supplementary Figure 2. Correlations within Vaccine Groups.** Heatmaps demonstrating all significant correlations between outcomes and spike antibody measurements when NHPs are analysed within their respective group (Two-sided Spearman correlation).

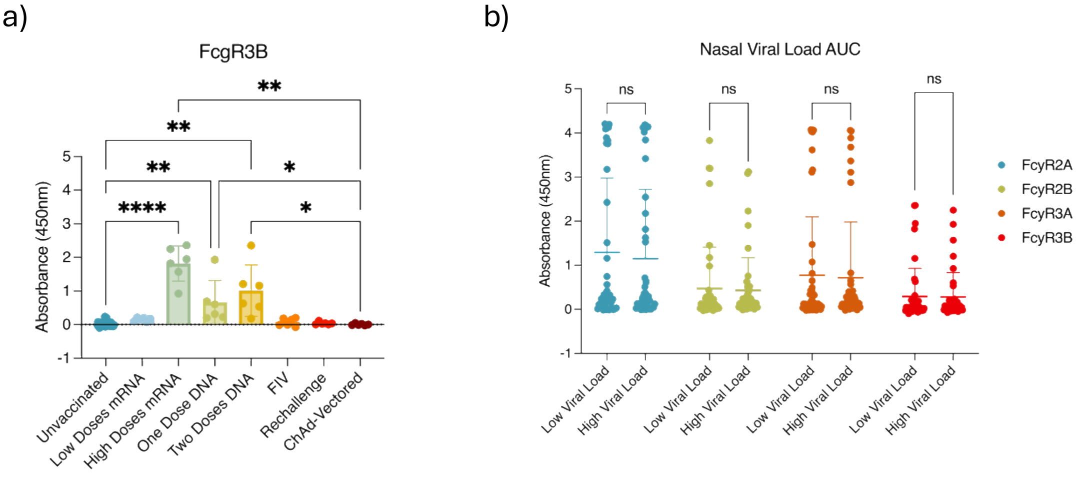


**Supplementary Figure 3. FcγR Binding. a)** FcγR3B binding by vaccine group. Dunn's multiple comparisons test performed. Datapoints represent individual macaques and the bars represent the mean and SD. **b)** FcγR binding comparison between nasal viral load AUC post-challenge groups. Tukey's multiple comparisons test performed. Datapoints represent individual macaques and the bars represent the mean and SD.

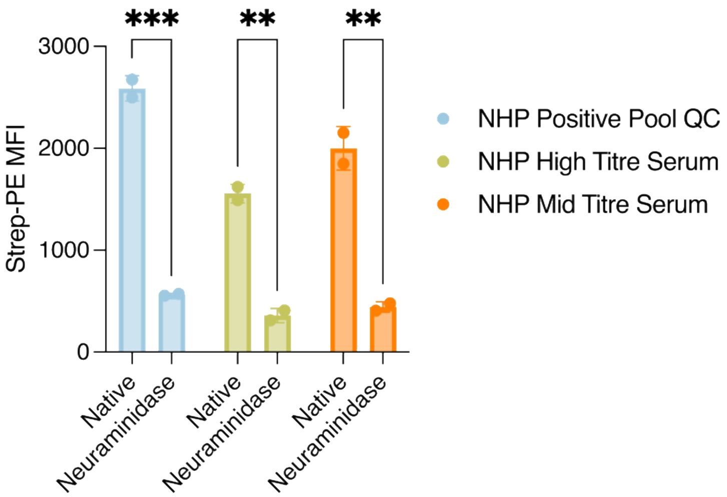


**Supplementary Figure 4. Sialic acid-specificity of SNA binding with neuraminidase-treated macaque serum.** Datapoints represent technical intra-assay replicates within the assay. Šídák's multiple comparisons test performed.

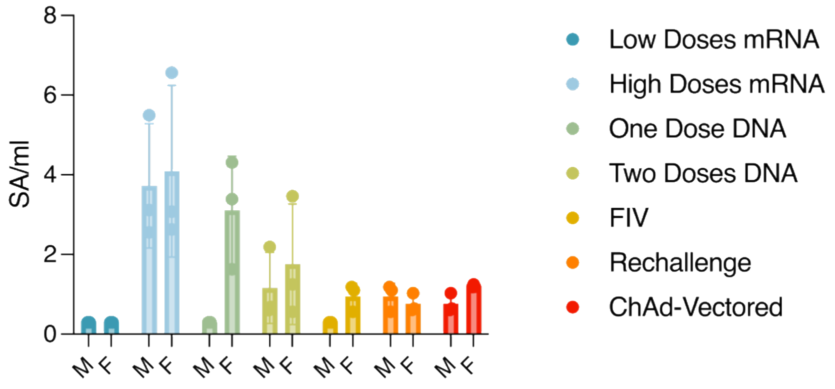


**Supplementary Figure 5. Spike antibody sialylation by sex in each immunisation group.** Two-way ANOVA and Tukey's multiple comparisons test performed. Datapoints represent each animal and the bars show group means with SD.

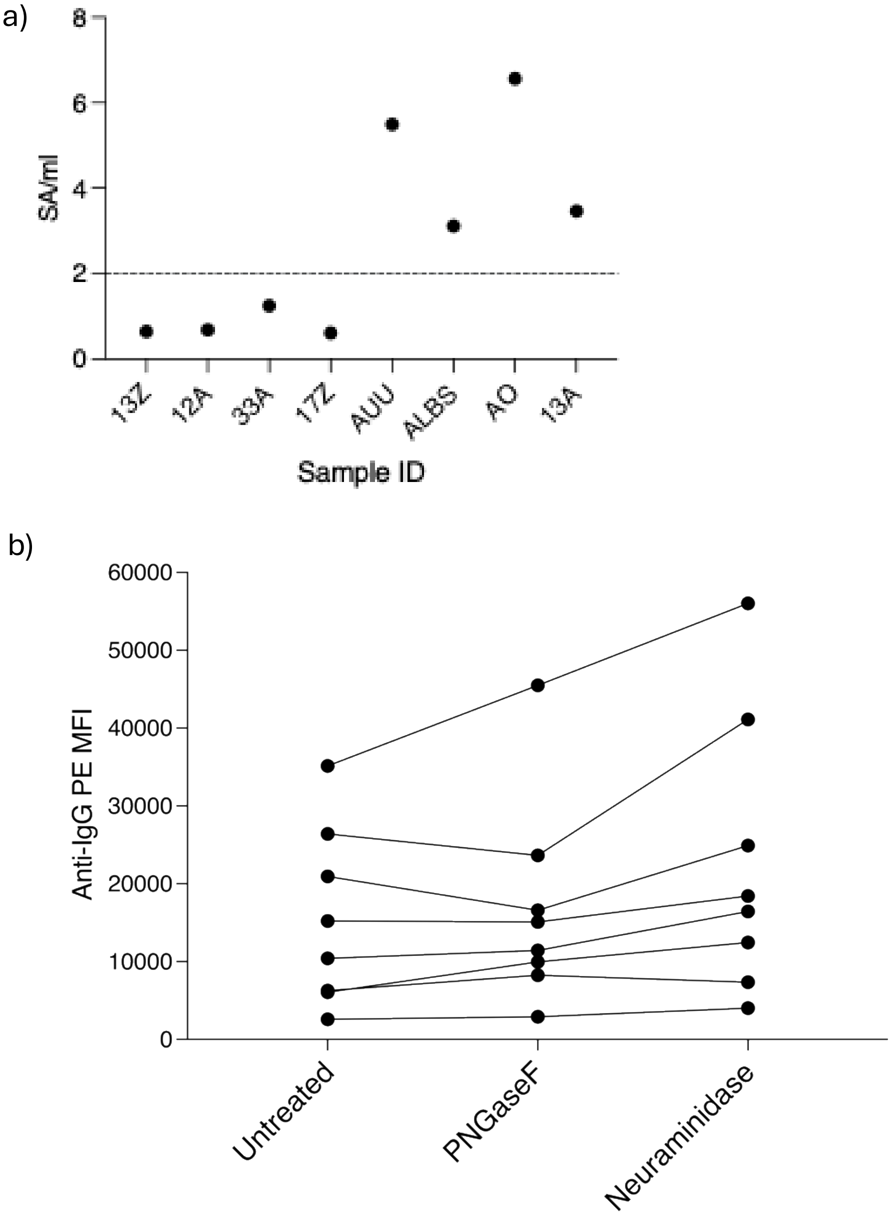


**Supplementary Figure 6. Glycosidase-treatment effect on spike bead binding of IgG. a)** The definition of the low sialylation versus the high sialylation group. **b)** Spike bead binding by native serum Ig versus serum treated with PNGaseF and neuraminidase. Datapoints represent the mean MFI of intra-assay technical replicates for each macaque sera tested. Wilcoxon paired t-test performed.

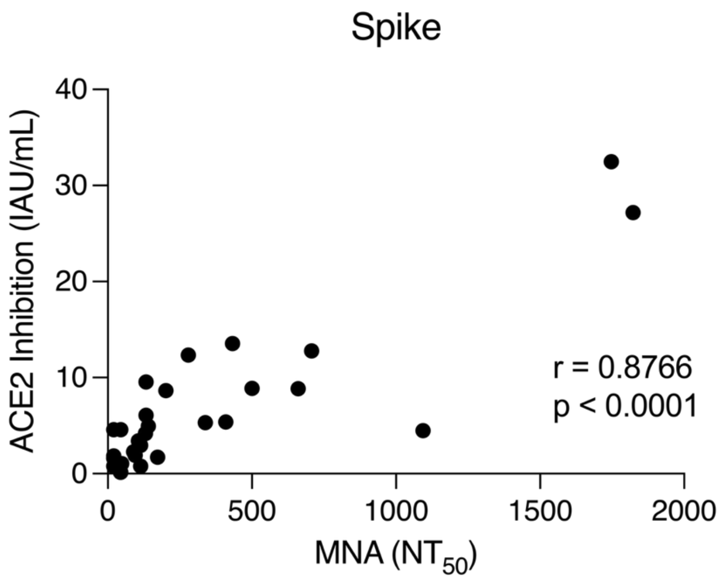


**Supplementary Figure 7. Correlation between ACE2 inhibition data and microneutralisation data.** Datapoints represent each macaque sera tested. Two-sided Pearson correlation performed.

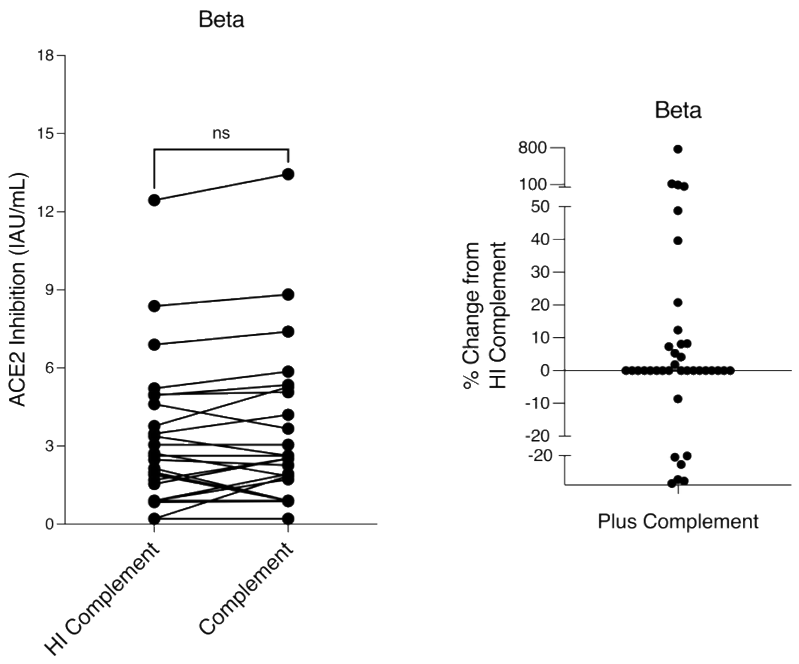


**Supplementary Figure 8. Complement-enhanced Beta ACE2 inhibition data data.** Datapoints represent each macaque sera tested, when heat-inactivated (HI) complement and non-HI complement is added to the ACE2 assay. Two-tailed paired student t-test performed. The percentage change in ACE2 inhibition when non-HI complement is added versus HI-complement is also demonstrated.

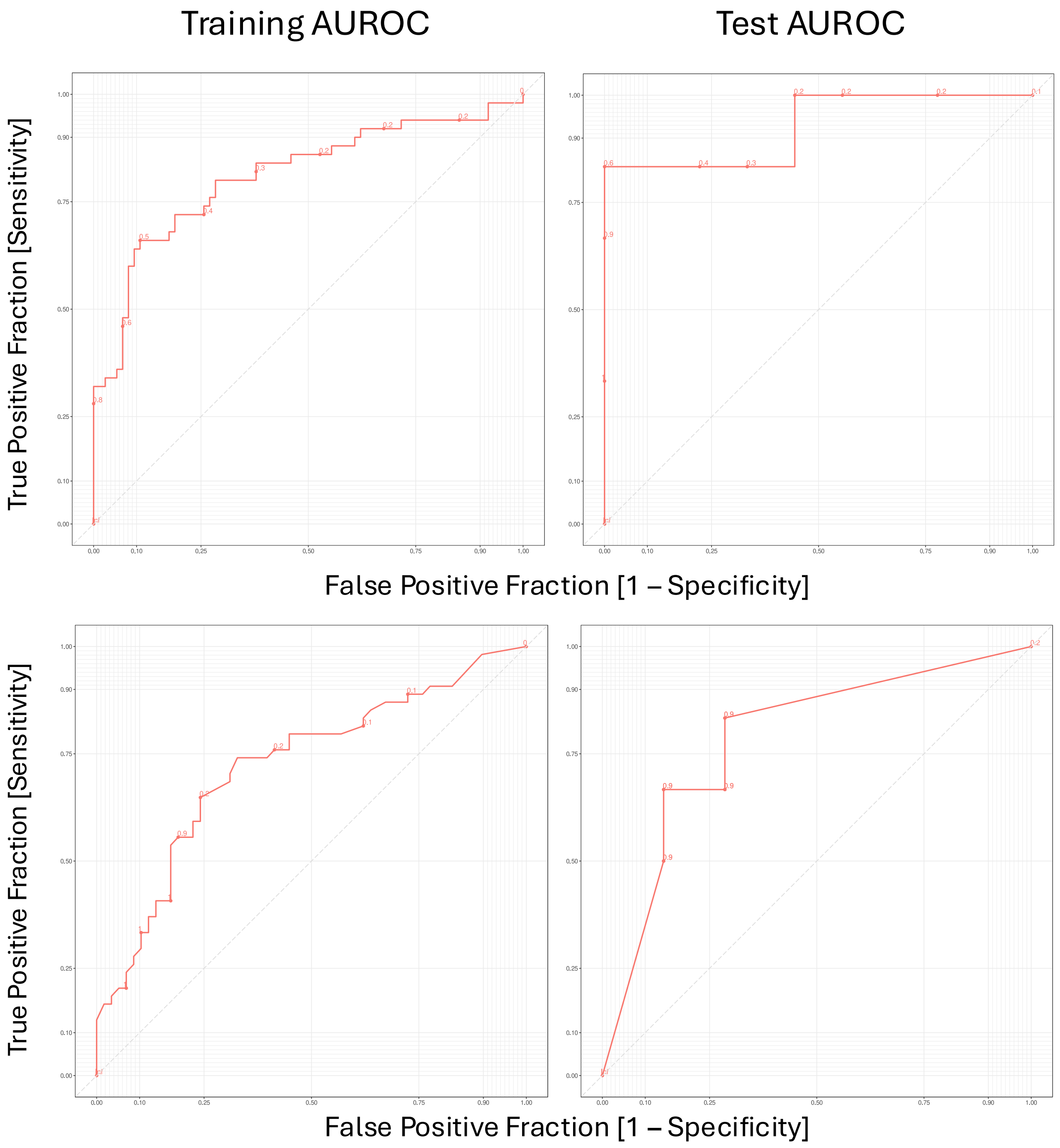


**Supplementary Figure 9. SIMON outputs for identifying the immune predictors of post-challenge outcomes.** The performance of the **a)** svmPoly model of clinical protection **b)** pcaNNet model of lung viral load 6-8 days post-challenge.

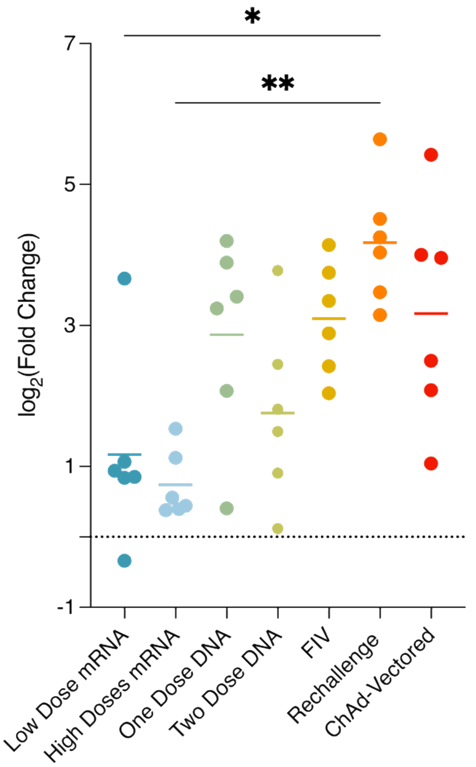


**Supplementary Figure 10. FcγR2A:FcγR2B binding ratio by vaccine group.** Log2(Fold Change) between FcγR2A and FcγR2B binding across groups. Dunn's multiple comparisons test performed. Datapoints represent individual macaques and the bars represent the mean.

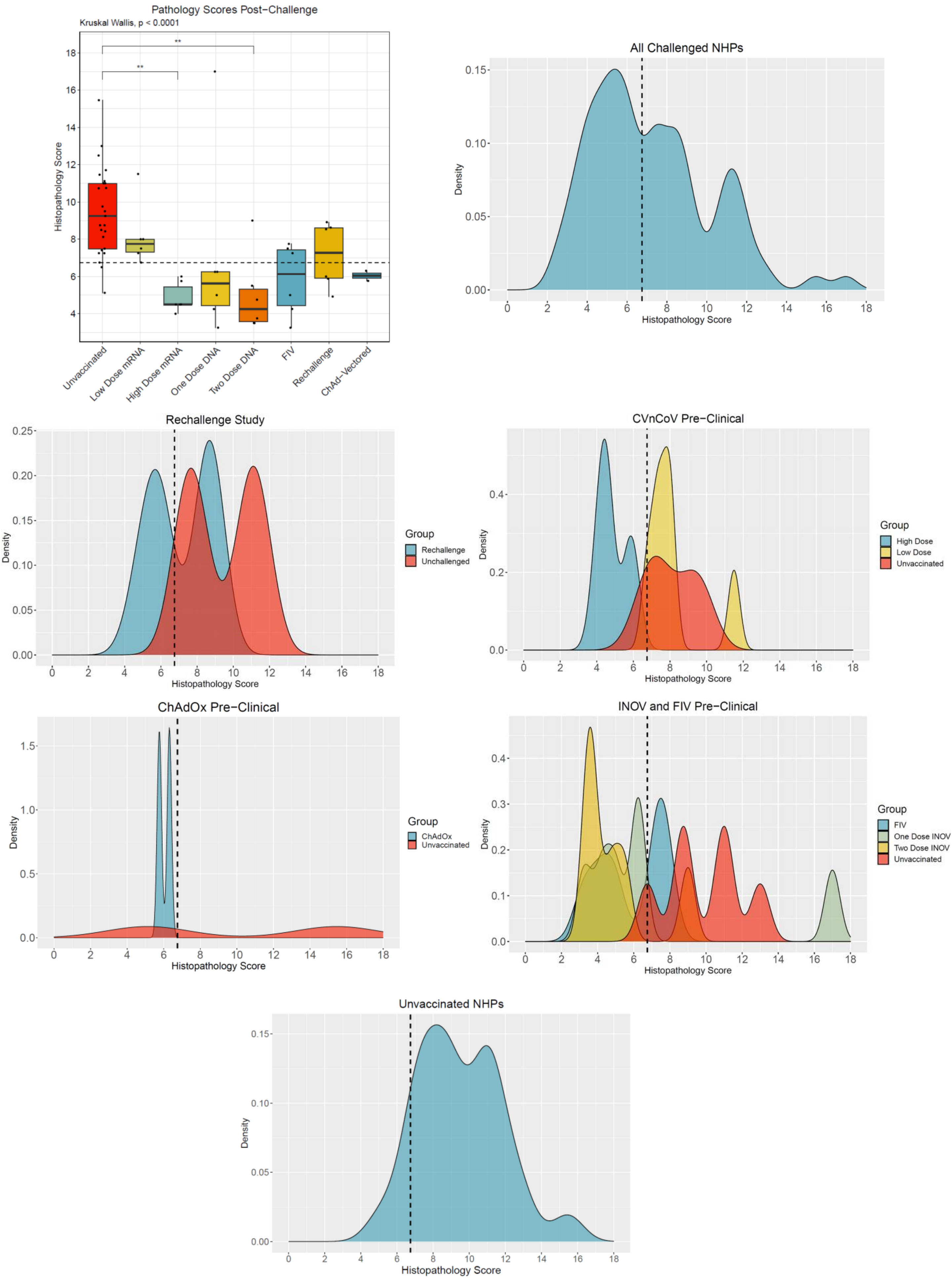


**Supplementary Figure 11. Defining the pathology outcomes.** Density plots demonstrate the spread of the data with the median used as the cut-off demonstrated in the bar blots.

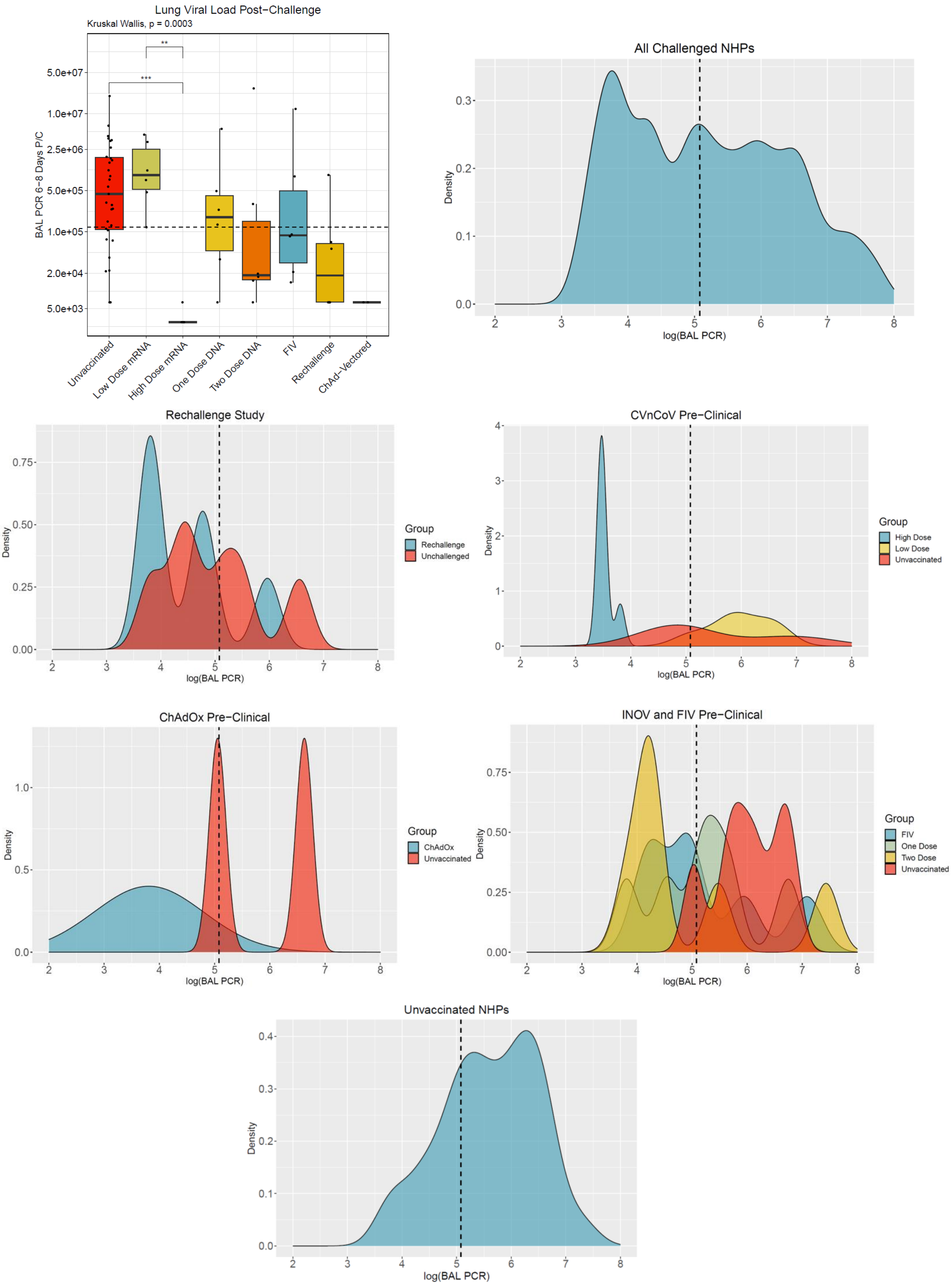


**Supplementary Figure 12. Defining the lung viral load outcomes.** Density plots demonstrate the spread of the data with the median used as the cut-off demonstrated in the bar blots.

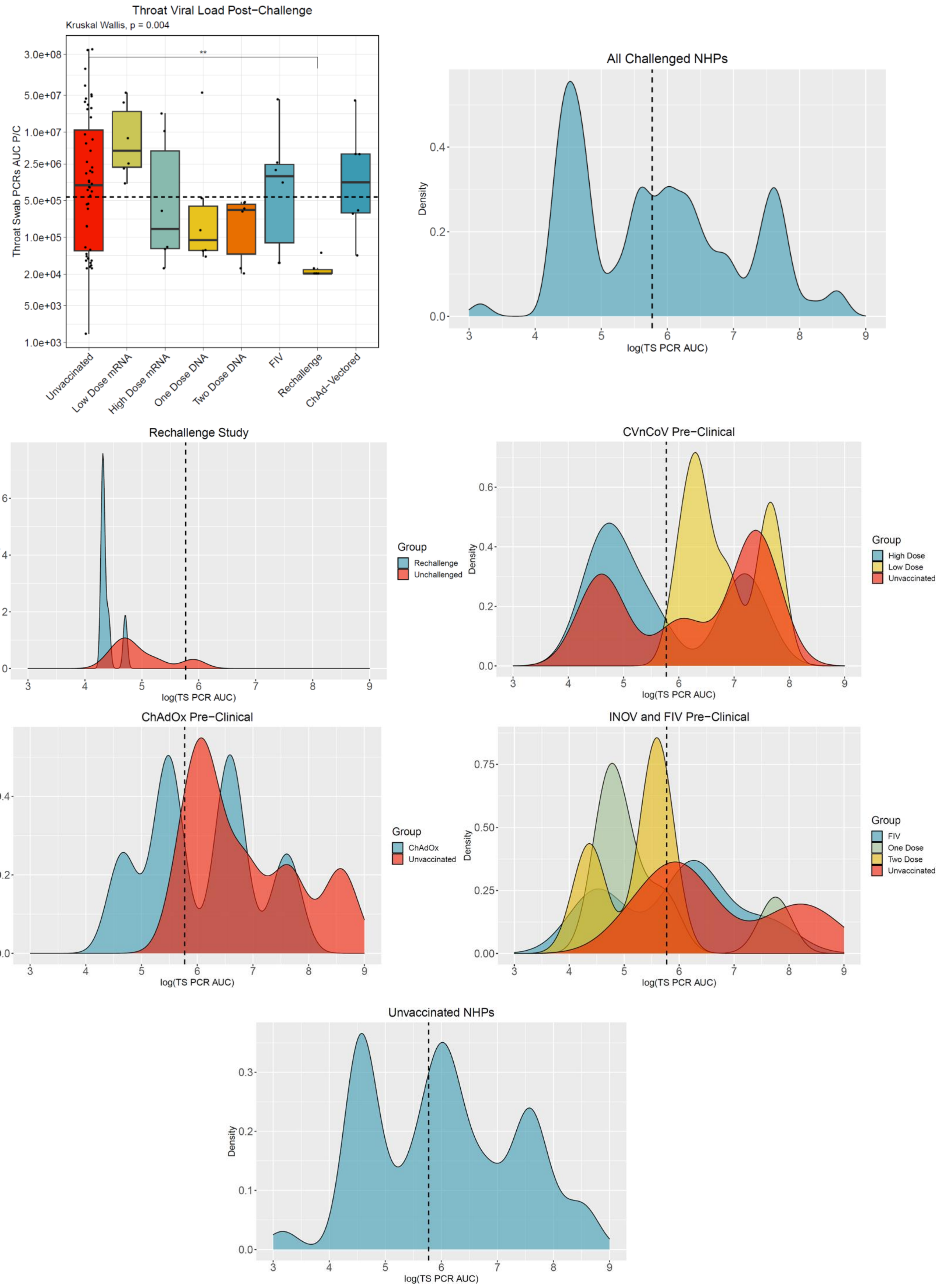


**Supplementary Figure 13. Defining the throat viral load outcomes.** Density plots demonstrate the spread of the data with the median used as the cut-off demonstrated in the bar blots.

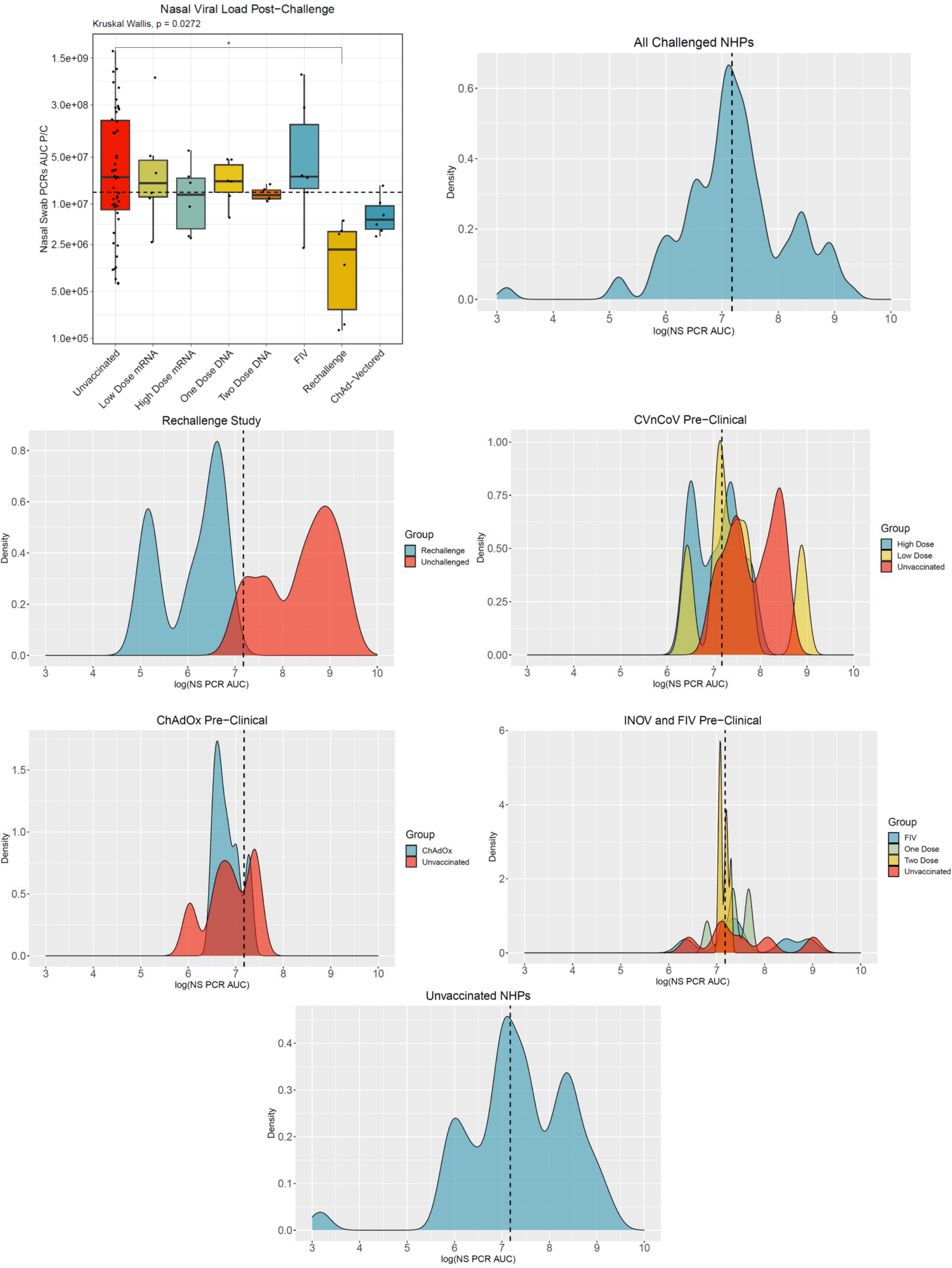


**Supplementary Figure 14. Defining the nasal viral load outcomes.** Density plots demonstrate the spread of the data with the median used as the cut-off demonstrated in the bar blots.
